# Structure-free, site-resolved contrastive learning extends small-molecule discovery beyond the reach of structure-based modeling

**DOI:** 10.64898/2026.07.28.741295

**Authors:** William E. Fondrie, Daniele Canzani, Lillian Tatka, J. Sebastian Paez, Anastasiya Prymolenna, Andrea Gutierrez, Julia Robbins, Brian McEllin, Evan Hubbard, Kyle Siebenthall, Lindsay K. Pino, Alexander J. Federation

## Abstract

Virtual screening asks which molecules, among an enormous space of drug-like chemistry, are worth synthesizing and testing against a protein target. Most modern methods answer this question by building and scoring an explicit three-dimensional pose through molecular docking, or the co-folding models that now approach experimental accuracy. Building these poses presumes a well-defined pocket. However, the non-orthosteric, cryptic, and intrinsically disordered sites where unexplored ligandability lies offer no such pocket to explore. Here we present Ptarmigan-1, a contrastive model that co-embeds the residues of a protein with candidate small molecules in a shared latent space, from sequence and two-dimensional chemistry alone, and without ever constructing a pose. Freed from the requirement for protein structures, Ptarmigan-1 trains directly on chemoproteomic and bioactivity data of mixed resolution, scores a compound in ten milliseconds rather than the tens of seconds a co-folding model demands, and resolves each prediction to the residues a compound engages. On well-folded, orthosteric targets it performs comparably to a collection of co-folding and docking models, and on covalent, cryptic, and disordered sites it matches or exceeds them. Ptarmigan-1 localizes reversible and covalent inhibitors to the pockets they engage, even for targets withheld from training, and screens the entire human proteome against a library of 3.4 billion compounds in under a day. By decoupling molecular recognition from structure, Ptarmigan-1 recasts virtual screening as a nearest-neighbor query in a rich latent space shared by protein residues and the compounds that bind them.

## Summary

- Ptarmigan-1 predicts which small molecules engage a protein directly from sequence and chemistry, with no three-dimensional pose, and screens a 3.4 billion compound library against the entire human proteome in under a day.
- It matches leading structure-based methods on standard, well-folded targets, and beats them on cryptic and poorly structured sites where those methods have no pocket to work from.
- It localizes engagement to the correct residues from sequence alone, generalizing to proteins and chemotypes withheld from training and discovering cryptic pockets.
- Precomputed embeddings make it thousands of times faster per target than a co-folding model, and the same embedding space answers proteome-wide queries as readily as single-target ones.
- Two patent-disclosed STAT6 inhibitor series, unseen in training, provide a retrospective test of the approach. Ptarmigan-1 recovers and localizes the actives of both to the SH2 domain, while a co-folding baseline mislocalizes the series whose binding site was never disclosed.

## Introduction

Despite our growing understanding of the molecular drivers of disease, few human proteins can be engaged with a small molecule. Of the roughly 20,000 human proteins, 704 are the target of an approved drug and 1,904 more have a potent small-molecule ligand, leaving 87% with neither [1, 2]. This is not a limitation of chemistry. The imaginable space of synthetically accessible, drug-like molecules is vast, estimated at anywhere from 10^33^ to 10^60^ structures [3, 4]. Any discovery campaign can synthesize or assay only an infinitesimally small slice of this space, so early discovery against novel targets is a problem of compound prioritization. Virtual screening attempts to address this computationally, searching large libraries *in silico* to nominate a tractable few. For most of its history, virtual screening has been tethered to protein structure. Molecular docking searches for the pose a compound adopts within a binding pocket and scores the resulting complex [5], and so cannot identify a site that was not nominated in advance. Co-folding models remove that requirement, predicting the three-dimensional (3D) structure of a protein–ligand complex directly, with AlphaFold [6, 7], Boltz [8, 9], and others [10–12] approaching experimental accuracy on many complexes. Both approaches, however, require the generation of a molecular pose itself, constructed for every protein–ligand pair at a cost of tens of seconds of compute. Docking has reached billion-compound libraries only by running for weeks across tens of thousands of processors [13, 14], and co-folding is more expensive still, placing the same library on the order of a thousand GPU-years against a single target.

Cost is not the only consequence of requiring a pose. Every prediction passes through a constructed complex; hence these models must be supervised on the coordinates of atoms in the structure and the binding affinity. Structureless affinity labels are usable, and co-folding models do train on them by conditioning an affinity head on predicted geometry [9, 15]. A measurement that names only the residue a compound engages, however, corresponds to nothing these models emit, and so cannot supervise them at all. The available data narrow the target space further. The complexes of the Protein Data Bank [16] and the affinities curated in ChEMBL [17], BindingDB [18], and PubChem [19] overwhelmingly describe well-folded proteins bound at their primary sites, and where coordinates are manufactured for structureless pairs they are retained only above a confidence threshold [15], which discards flexible and disordered targets first. On targets whose ligands lie outside orthosteric pockets, these methods tend to misplace known binders into the orthosteric pocket, an outcome the authors termed “orthostery burnout” [20]. For non-covalent ligands, a panel of AlphaFold3, Protenix, Boltz-2, Chai-1, and DynamicBind struggles on allosteric complexes relative to orthosteric ones [21], and covalent and site-specific predictions show the same shortfall [22]. Benchmarks built to test whether these models memorize rather than generalize [23] have driven substantial gains on the least-precedented cases [24], but every prediction still flows through a biased predicted pose.

Much of the undrugged proteome does not resemble the well-folded proteins in these databases. For example, transcription factors act through extended, shifting surfaces rather than compact pockets, which is why they have resisted small-molecule development despite decades of interest [25]. Over 200 transcription factors have been observed as dependencies across cancer [26] and roughly a fifth of cancer-driver mutations fall within intrinsically disordered regions [27]. Ligands for exactly these proteins are already being found experimentally. Activity-based protein profiling (ABPP) maps reactive, ligandable sites directly in the native proteome [28, 29], and has assigned ligands to hundreds of proteins bearing sites far from any canonical pocket [30, 31], among them targets long deemed undruggable, such as transcription factors. These experiments report the residue a compound modifies rather than the geometry of a complex. A model that requires a pose cannot train on them, so the measurements that describe the undrugged 87% are unavailable to the methods requiring a 3D pose.

Here we present Ptarmigan-1, a contrastive model that co-embeds the residues of a protein with candidate small molecules in a shared latent space and never builds a pose. Unlike previous models that compact protein representations into a single vector [32–34], or that leverage cross-attention between the protein and ligand [35, 36], Ptarmigan-1 embeds every residue of the protein sequence and the putative ligand into a latent space where distance of a ligand to a residue indicates its likelihood of engagement. In turn, scoring a compound costs only milliseconds, which makes it possible to screen billions of compounds across the entire proteome without an extensive compute budget. This formulation also provides residue-resolved binding predictions without the expensive reintroduction of structure or pocket rendering used by other methods [32, 37, 38]. Ptarmigan-1 trains directly on the chemoproteomic and bioactivity measurements that structure-based models cannot use, which is how it reaches the flexible, disordered, and non-orthosteric targets that such models have not observed. Furthermore, Ptarmigan-1 scores every residue, predicting not only whether a compound will bind, but also where it will engage the target of interest—even on disordered regions that present no previously known pocket.

## Results

### Ptarmigan-1 is a model for fast and efficient virtual screening across the proteome

Ptarmigan-1 is designed to predict protein–ligand molecular engagement without resolving an explicit three-dimensional structure (Figure 1A). The model couples two foundation models, ESM Cambrian for protein sequences [40] and ChemBERTa for compound SMILES [41], each adapted to the task by parameter-efficient fine-tuning (PEFT) of low-rank adapters rather than by full retraining. Neither foundation model was trained on protein–ligand complexes, so neither inherits the bias toward crystallizable proteins and well-characterized pockets. Ptarmigan-1 instead learns engagement from protein sequence and chemistry alone. We trained the model with two paired contrastive objectives, one that operates over individual residues, and one over the whole protein, allowing the single model to learn from data sources of varying levels of resolution (Supplementary Figure 1). The residue-level objective draws each compound toward the residues it engages, while the protein-level objective requires only a binary record of whether a compound engages a protein at all.

**Figure 1:**
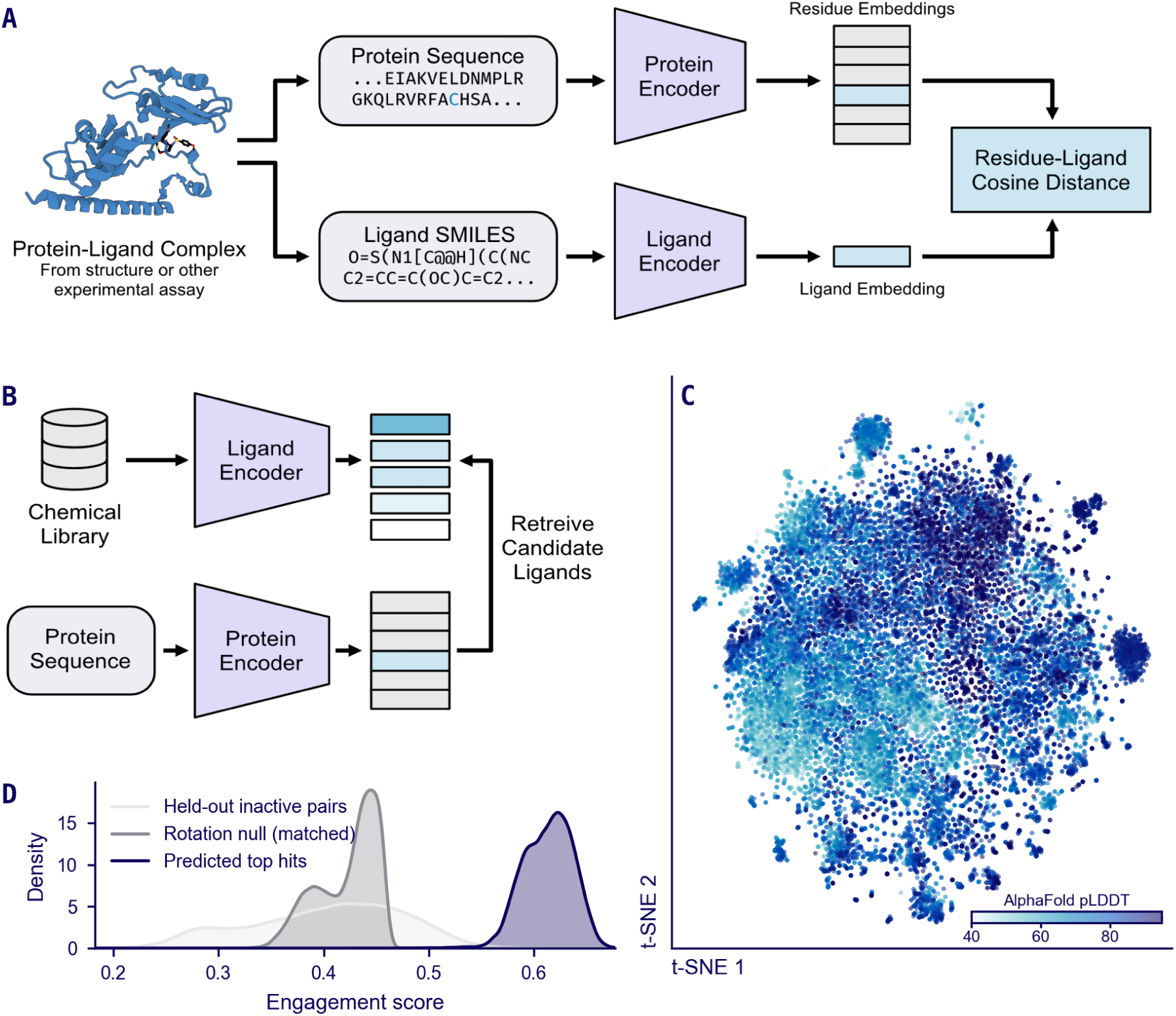
Model architecture, retrieval by precomputed embeddings, proteome coverage, and proteome-wide top-hit engagement. (A) Ptarmigan-1 encodes a protein sequence and a ligand SMILES string with two fine-tuned foundation models and embeds each residue near the ligands that engage it. Engagement is the residue–ligand cosine similarity, and a temperature-scaled softmax over residues yields a protein-level score. (B) Engagement compares precomputed embeddings and an approximate nearest-neighbor index retrieves the compounds with embeddings falling nearest to the residues of a target. (C) A t-SNE of all human proteins screened by Ptarmigan-1, positioned by the mean of each protein’s residue embeddings and colored by AlphaFold-predicted fold confidence (mean pLDDT). (D) Engagement-score distributions from screening every human protein against the full OnePot CORE library [39] (3.4 billion compounds). The best-compound engagement for each protein (dark) is compared against a matched rotation null (grey): the same maximum-over-library selection after a random rotation of the residue embeddings destroys the learned protein–compound alignment while holding the library and embedding geometry fixed (Methods). A naive, unmatched reference (light) shows the distribution of held-out inactive protein–ligand pairs, which are individual pairs rather than a maximum.

Ptarmigan-1 scores a residue–ligand pair by the proximity of their embeddings. To screen a library against a target, we embed the compound library once and then retrieve the ligands whose embeddings fall nearest the target residues (Figure 1B). An approximate nearest-neighbor index keeps this retrieval fast even for libraries of billions of compounds, and the embedded library is reused for every subsequent target. To turn per-residue predictions into a protein-level prediction, Ptarmigan-1 aggregates the residue–compound similarities through a temperature-scaled softmax, so that a small number of strongly engaged residues determine whether the protein is scored as engaged. Since the protein-level prediction needs no residue-level label, Ptarmigan-1 can train on assays that report only a binding outcome alongside those that localize binding to the residue-level.

With the computationally trivial cost of inference using this architecture, the entire human proteome can be scored against large chemical space, here a 3.4 billion compound library. Retrieving the top predicted ligands for every protein took under a day (20 H100 GPU-hours). The screen itself spanned the 20,431 proteins of the human proteome with a range of confidence in their predicted structures, and included even the most intrinsically disordered proteins in the proteome (Figure 1C). That breadth let us ask, across the entire proteome, how strongly the best compound in a large library engages each protein. For virtually every protein, the single best-scoring compound scored far above a matched rotation null (Figure 1D, Methods), a re-screen against the same library in which a random rotation of the residue embeddings destroys the learned protein–compound alignment while preserving the embedding geometry. The real top-hits’ 10th percentile engagement logit (0.33) alone exceeds the rotation null’s maximum (−0.11). On a stratified subsample of 1,000 proteins re-screened under 40 independently drawn rotations, every protein exceeded all 40 of its own null draws (empirical *p* = 0.024, the smallest value 40 draws permit; Benjamini–Hochberg *q* = 0.024 across the subsample), including all 177 predicted to be intrinsically disordered (AlphaFold mean pLDDT below 50). Because the rotation scrambles only the residues’ orientation toward the compounds and leaves the protein representation otherwise intact, this gap reflects learned engagement rather than the inevitability of a high maximum over such a vast library. We therefore read the panel as suggestive that protein–ligand engagement may extend universally across the proteome, including its most disordered members.

### Ptarmigan-1 performs competitively on a standard structured benchmark

We first assessed Ptarmigan-1 on well-folded, structured targets that co-folding and docking methods handle best. We intentionally held LIT-PCBA targets out of the training data for Ptarmigan-1, then compared the baseline performance of co-folding, docking, and contrastive models. When possible, previously published results for the LIT-PCBA targets were used [42]. An important caveat for this benchmark is that the training data for Boltz-2 and other methods may include examples from the LIT-PCBA active targets.

We measured ranking quality with the adjusted logAUC, where a random ranker scores 0 and a perfect ranker approaches 0.855, scoring an identical set of compounds under every method. The co-folding model Boltz-2 led on average across these five targets, with Ptarmigan-1 ranking second, above the other docking, co-folding, and co-embedding methods with available scores. Performance remained competitive on the early-recognition metrics EF@1% and BEDROC (Table 1). A breakdown of LIT-PCBA active recognition by ligand novelty, for Ptarmigan-1 with and without internal training data and the published baselines, is given in Supplementary Figure 2.

**Table 1:** LIT-PCBA leaderboard, averaged over the five target-sets with published structure-based baselines. The last two columns give each method’s class and whether it runs structure-free: docking (Glide-SP, Gnina-Vina, Gnina-CNN) and co-folding (Boltz-2, Protenix) construct an explicit three-dimensional pose, whereas the co-embedding methods (Ptarmigan-1, ConPLex, SPRINT) score a protein–ligand pair by proximity in a shared latent space. Of these, only Ptarmigan-1 and ConPLex [33] are structure-free, scoring from protein sequence and two-dimensional chemistry alone; SPRINT [34], though also a co-embedding method, requires a three-dimensional structure, which it encodes as a SaProt fold. Every method scored an identical set of compounds, each score oriented so that higher indicates greater predicted activity (Methods). Metrics are ROC-AUC, adjusted logAUC (*λ* = 0.001, random = 0), enrichment factor and normalized enrichment factor at 1% (EF@1%, NEF@1%), and BEDROC (*α* = 20); rows are sorted by adjusted logAUC.

| model | ROC-AUC | logAUC | EF@1% | NEF@1% | BEDROC | Method | Structure-free |
| --- | --- | --- | --- | --- | --- | --- | --- |
| Boltz-2 | 0.776 | 0.196 | 9.75 | 0.116 | 0.264 | Co-folding | No |
| Ptarmigan-1 | 0.672 | 0.12 | 7.32 | 0.084 | 0.183 | Co-embedding | Yes |
| Glide-SP | 0.642 | 0.11 | 6.79 | 0.078 | 0.172 | Docking | No |
| Protenix | 0.61 | 0.091 | 7.58 | 0.088 | 0.153 | Co-folding | No |
| SPRINT | 0.694 | 0.091 | 1.79 | 0.028 | 0.124 | Co-embedding | No |
| ConPlex | 0.659 | 0.082 | 4.32 | 0.047 | 0.117 | Co-embedding | Yes |
| Gnina-Vina | 0.652 | 0.073 | 2.82 | 0.038 | 0.106 | Docking | No |
| Gnina-CNN | 0.576 | 0.071 | 6.1 | 0.067 | 0.143 | Docking | No |

Direct comparison to DrugCLIP [32] was not possible within the context of Table 1 and is reported in Supplementary Table 2, restricted to the two quantities whose definition is independent of its evaluation pipeline. DrugCLIP co-embeds protein and ligand like Ptarmigan-1 but encodes the protein as a three-dimensional Uni-Mol binding pocket rather than from sequence, and so requires a resolved pocket. Its published performance comes from its own evaluation pipeline with its own metric set (BEDROC at *α* = 80.5 rather than the *α* = 20 used throughout, and no adjusted logAUC).

### Ptarmigan-1 localizes ligands to the residues they engage

Ptarmigan-1 embeds each residue of a protein and each compound into a shared latent space and scores the engagement between a residue and a compound as the cosine similarity of their embeddings, without constructing a pose. Scoring one compound against every residue of a target therefore returns a residue-level map of engagement, showing where along the sequence the compound is predicted to bind rather than only whether it binds. We asked (1) whether these maps are accurate, (2) whether their accuracy holds for proteins and chemotypes withheld from training, and (3) whether it depends on our internal training data.

Covalent inhibitors are a stringent test of localization, because the residue each one modifies is known exactly. We scored four covalent inhibitor–target pairs, each with a co-crystal structure that defines the reactive cysteine, from protein sequence and compound SMILES alone (Figure 2A). For each pair, engagement peaked sharply at the reactive cysteine, along with at other residues in the annotated pocket. BTK/ibrutinib was in the Ptarmigan-1 training data, whereas BMX/ibrutinib (a known ibrutinib off-target), the KRAS G12C mutant/ adagrasib, and ITK/PRN694 were held out. Coloring residues with the BTK binding pocket by predicted engagement score shows the signal concentrated on the residues that contact the bound inhibitor (Figure 2B).

**Figure 2:**
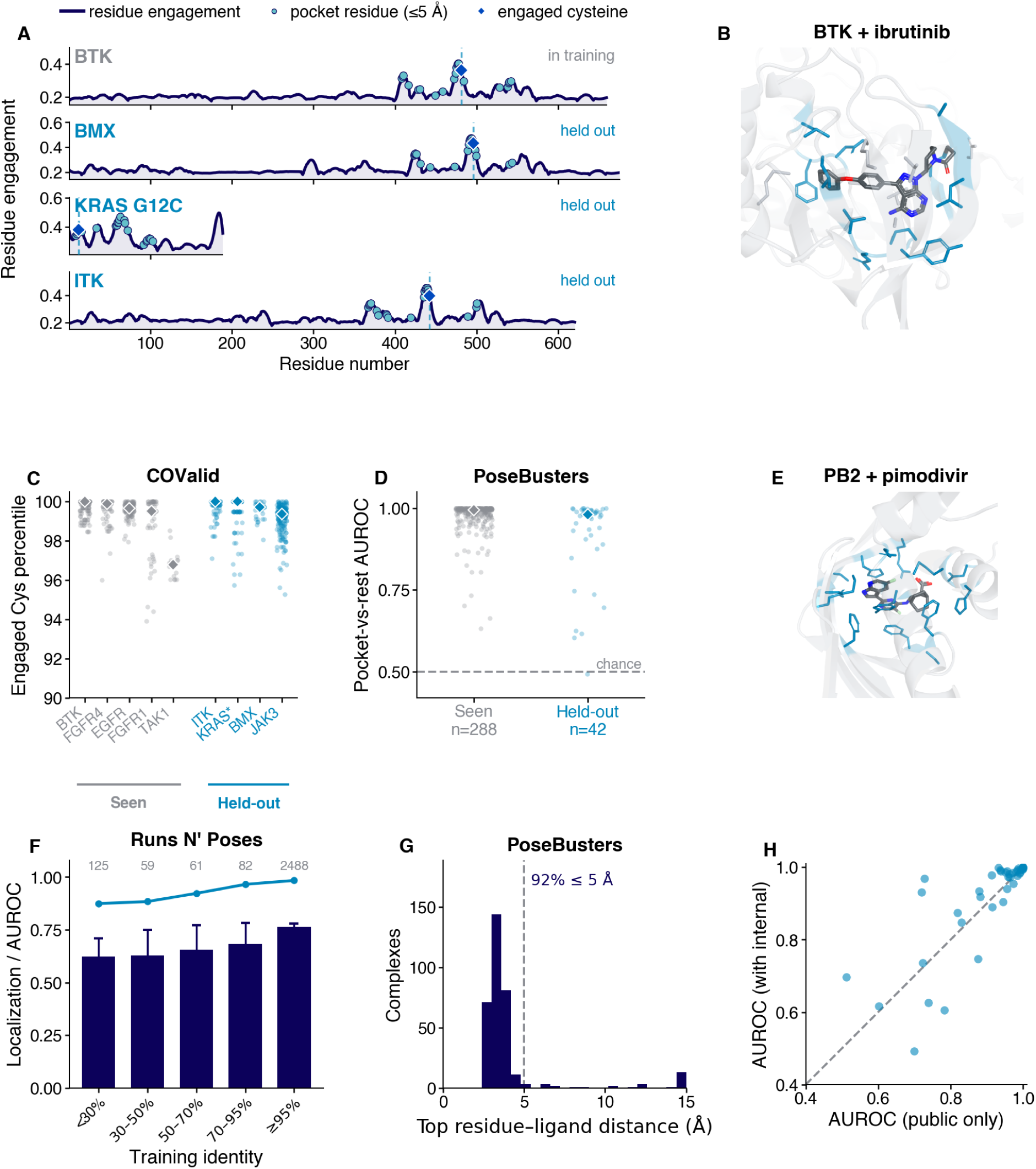
Residue-level localization of covalent and non-covalent ligands, for targets seen in and withheld from training. (A) Full-length smoothed per-residue engagement for four covalent inhibitor–target pairs: BTK/ibrutinib (in training) and three pairs held out of training, BMX/ibrutinib (a known ibrutinib off-target), the KRAS G12C mutant/adagrasib, and ITK/ PRN694. Turquoise circles mark the pocket residues (within 5 Å of the co-crystallized ligand in the reference structure: BTK, PDB 5P9J; BMX, 8X2A; KRAS G12C, 6OIM; ITK, 9NWX); the cobalt diamond and dashed line mark the engaged cysteine. (B) The BTK/ibrutinib pocket (PDB 5P9J) with pocket residues drawn as sticks and two-toned by engagement (grey below, sea-blue above 0.26), with ibrutinib bound in the cleft. (C) Engaged-cysteine percentile across the COValid benchmark [43] containing 874 covalent actives across nine targets, each with an experimentally determined engaged cysteine. Each point is one active’s engaged-cysteine percentile among all residues; diamonds mark per-target medians; targets are grouped and colored as seen in training (grey) or held out (blue). KRAS* denotes the KRAS G12C mutant. (D) Pocket-vs-rest AUROC across the PoseBusters set [44] of non-covalent, drug-like complexes: for each complex, how well per-residue engagement separates the crystallographic pocket (within 5 Å of the ligand) from the rest of the protein. Points are complexes with a target seen in training (*n* = 288) versus held out (*n* = 42); diamonds mark the group medians (0.99 and 0.98) and the dashed line marks chance (0.5). (E) The influenza PB2 cap-binding domain with pimodivir (PDB 7AS1), held out of training, rendered as in (B) with the sea-blue threshold at 0.30. (F) Localization on the Runs N’ Poses benchmark [23], binned by sequence identity to the nearest training protein. Bars, fraction of systems whose top-engagement residue lies within 5 Å of the annotated ligand site (exact-binomial 95% CI); line, median pocket-vs-rest AUROC; per-bin counts above. The companion analysis by ligand novelty is in Supplementary Figure 3. (G) Distribution of the distance from Ptarmigan-1’s single top-engagement residue to the ligand, across drug-like PoseBusters complexes (*n* = 337), from sequence alone. The dashed line marks the 5 Å binding-site threshold, our residue-level analog to the pocket-recovery criterion structure-based tools report [45], labeled with the resulting success rate (92%). Co-folding and docking methods report a different quantity, pocket-aligned pose RMSD < 2 Å, and are compared in the main text rather than plotted here. (H) PoseBusters per-complex pocket-vs-rest AUROC for held-out drug-like complexes (*n* = 42), trained on public data only (*x*) versus with the internal corpus added (*y*); dashed line, identity.

To extend the evaluation of localization accuracy of Ptarmigan-1 at scale, we scored every covalent active in the COValid benchmark [43], 874 compound–target pairs across nine targets, each with an experimentally determined engaged cysteine (Figure 2C). The engaged cysteine fell within the top 1% of residues for eight of the nine targets (median percentile ≥ 99), with the exception of TAK1, which had the fewest actives and sat near the 97th percentile. Four of these targets were withheld from training (JAK3, the KRAS G12C mutant, BMX, and ITK), and the model localized them as precisely as the five it had seen in training.

Localization was not specific to covalent chemistry. On the PoseBusters set [44], a benchmark of non-covalent, drug-like complexes, we scored each residue’s engagement with the crystallized ligand without building a pose, then used the deposited structure to label the residues within 5 Å of that ligand as its pocket and, per complex, took the AUROC for ranking those pocket residues above the other residues by engagement (Figure 2D). This pocket-vs-rest AUROC had a median of 0.99 across the benchmark. Pocket identification held for proteins whose sequence was absent from training (median 0.98, *n* = 42). As an example, we rendered the cap-binding domain of influenza PB2 bound to pimodivir, an interaction held out of training (Figure 2E).

We next turned to Runs N’ Poses [23], a more recent benchmark of 2,815 ligand systems built to probe memorization vs. generalization for co-folding models. Because Ptarmigan-1 learns from binding assays and not from PDB structures alone, it is not bound by the training cutoff that Runs N’ Poses assumes. About 96% of these proteins (≥ 30% identity to a training protein) and 98% of these ligands (≥ 0.4 Tanimoto to a training ligand) fall above the novelty cutoffs the benchmark uses, so its similarity axis does not measure novelty for our model. We instead re-binned each system by sequence identity to the nearest protein in Ptarmigan-1’s training set. Localization fell only modestly with novelty. For proteins below 30% identity to proteins in training (*n* = 125), the top-engagement residue still reached the ligand in 62% of systems at a median pocket-vs-rest AUROC of 0.87, rising to 76% and 0.98 for proteins seen in training (Figure 2F), and was similarly robust to ligand novelty (Supplementary Figure 3). This gradient is the expected signature of generalization rather than recall, with performance highest on familiar proteins and well above chance even on those with no training homolog.

We define binding-site recovery, our residue-level analog to the pocket-recovery criterion structure-based tools report [45], as whether the single top-scoring residue of a complex lies within 5 Å of the ligand. By that criterion, Ptarmigan-1 succeeded for 92% of drug-like complexes from sequence alone (Figure 2G). For comparison, given the structure or pocket, AutoDock Vina [5] and DiffDock [46] placed a full, physically valid pose (RMSD < 2 Å) for 53% and 40% of these same 337 drug-like complexes, recomputed by us on that subset. The published blind co-folding success rates for AlphaFold 3 [7] and Boltz-2 [9], 76% and 77%, are reported over all 428 complexes of the PoseBusters set rather than this subset, so they share neither our denominator nor our outcome measure. Recovering the binding site is a weaker requirement than reproducing a full pose, so this is not a head-to-head comparison. Rather, it does show that Ptarmigan-1 surfaced the true binding site from sequence alone at least as often as these methods reproduce the pose from a given structure.

Pocket recovery alone cannot show that a prediction is specific to the compound supplied, since a model that had learned only where each protein is ligandable would place its top residues in the same pocket. To test this, we rescored each complex against a property-matched ligand from a different protein and observed a loss in binding-site recovery from 92% to 77% (paired, *p* < 10^−14^; Supplementary Table 1), suggesting that localization is conditioned on the query compound rather than reporting a fixed ligandable pocket.

Finally, we asked whether this localization depended on our internal training data. Retraining on public data alone and re-scoring the held-out PoseBusters proteins leŁ the per-complex AUROC essentially unchanged (median 0.96, versus 0.98 with the internal corpus added), with the points falling along the identity line (Figure 2H).

Localization is thus a property of the sequence-based contrastive architecture, reproducible from public data alone, and not an artifact of the internal corpus.

### Ptarmigan-1 screens orders of magnitude faster and compares targets in a shared embedding space

Because Ptarmigan-1 scores a compound–protein pair by the proximity of precomputed embeddings rather than by constructing a pose, it screens at a throughput much higher than pose-based methods. Embedding a library is a one-time cost that grows approximately linearly with its size. Once embedded, retrieval against an approximate nearest-neighbor index stayed between 10 and 40 seconds for libraries from one thousand to ten million compounds, and every additional target reused the same index (Figure 3A, Figure 3B). Per compound, Ptarmigan-1 scored a protein–ligand pair in 10 ms against 54 s for Boltz-2, a roughly 5,000-fold speedup that shortens a million-compound screen from nearly two years to under three hours on a single GPU (Figure 3C).

**Figure 3:**
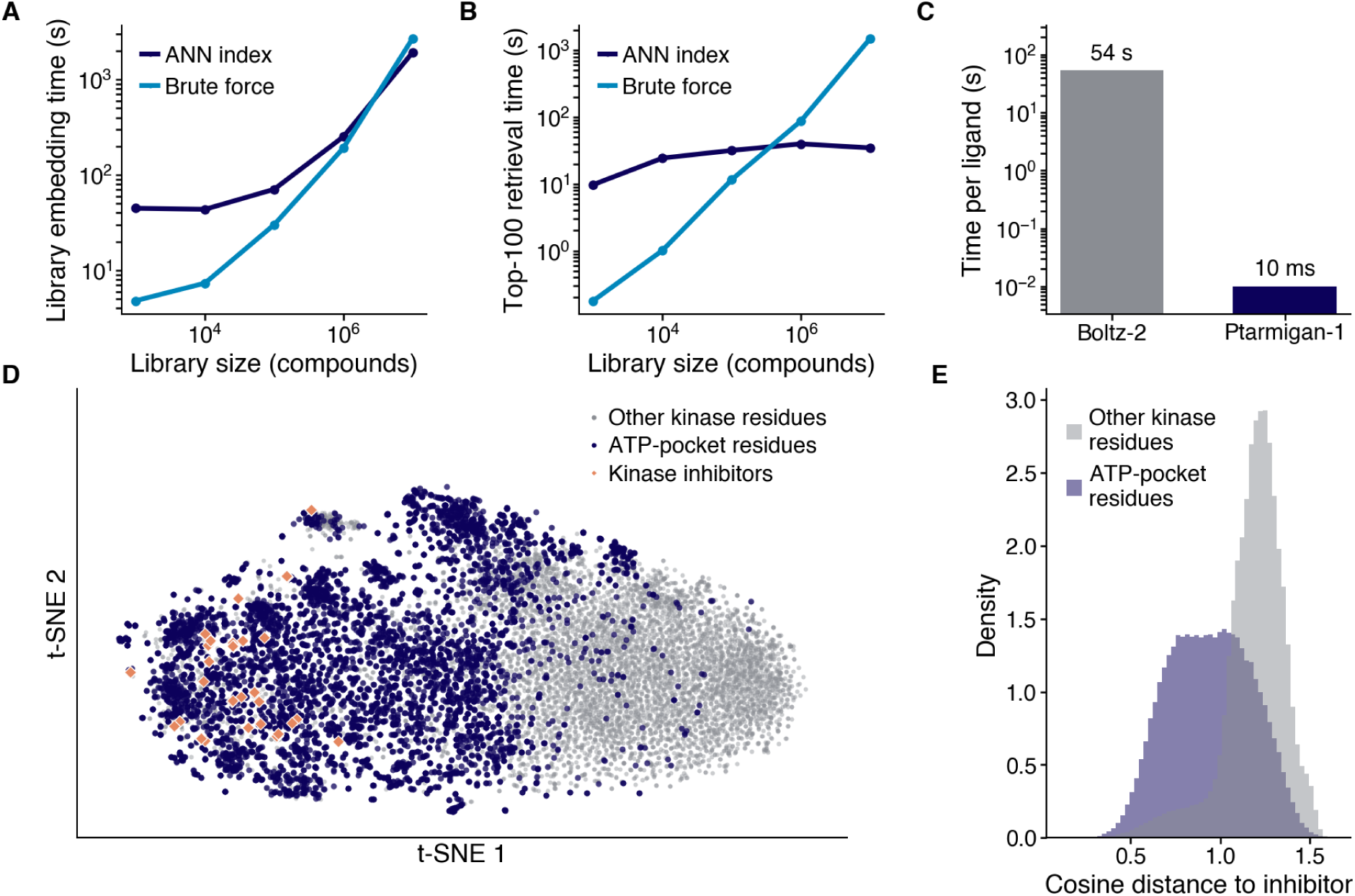
Library embedding and retrieval times, per-compound scoring time, and the shared embedding space across 560 kinases. (A) Library embedding time as a function of library size (log–log; one H100 GPU) for brute-force in-memory encoding and building an approximate nearest-neighbor (ANN) index. (B) Time to retrieve the top-100 compounds for a target as a function of library size (log–log). (C) Per-ligand time from SMILES to score for a single target (EGFR; log scale), including compound embedding: 10 ms for Ptarmigan-1 and 54 s for Boltz-2. (D) A joint t-SNE of protein residues and compounds in the shared embedding space. ATP-pocket residues of 560 human kinases are colored blue along with a seeded sample of the remaining residues of those same proteins. 46 approved kinase inhibitors shown co-embedded as diamonds. Every embedding in panels D and E was computed in 42 s of inference on a single H100 (40 s for the 395,403 residues of the 560 kinases and 1.4 s for the inhibitors). (E) Cosine distance from each inhibitor to the ATP-pocket residues and to a seeded sample of the other residues of the same kinases. Median cosine distance to the pocket is 0.94 versus 1.21 to the other residues.

This throughput follows from the shared embedding space between a ligand and the residues it engages, and that same space supports the inverse query. Rather than ranking many ligands against one protein, we can rank many proteins against one ligand. To illustrate, we embedded the residues of 560 human kinases alongside 46 FDA-approved kinase inhibitors (Figure 3D). The inhibitors lay closer to the ATP-pocket residues than to the other residues of the same kinases (median cosine distance 0.94 versus 1.21), so a single embedding captures both which compounds engage a protein and where they bind, across many proteins at once (Figure 3E). Computing every embedding for this analysis took roughly 42 seconds on a single GPU.

### Ptarmigan-1 generalizes to targets and chemotypes withheld from training

Fast screens against large chemical space are only useful if compound rankings generalize to proteins and chemistry the model has not yet seen. To probe generalization, we assessed our internal held-out screening benchmark (100 targets drawn from the test splits of our public sources, 33 of them proteins whose sequence never appears in training; Supplementary Figure 4, Methods). Ptarmigan-1 enriched actives above random for 32 of the 33 held-out targets, at a median adjusted logAUC of 0.31.

This enrichment climbed with training-protein coverage (from a median adjusted logAUC of 0.16 for targets with few distant homologs to 0.36 for those with dozens, Figure 4A). However, even the least-precedented targets enriched actives well above random. Targets whose sequence appeared in training scored higher still (median 0.65, Supplementary Figure 4), the upper reference line in Figure 4A, following the expected benefit of prior exposure. The held-out targets carry no target-specific training, so their enrichment from protein sequence and two-dimensional chemistry alone implies that Ptarmigan-1 does not require previous exposure to rank actives highly. We then asked whether this enrichment reflects genuine molecular recognition, or is reporting generic drug-likeness or leakage of training analogs into the benchmark. Binning every test-set active by its ECFP4 Tanimoto to the nearest training compound, the pooled adjusted logAUC rose with chemical familiarity, from 0.21 for the most novel ligands to 0.43 for the most precedented (Figure 4B). Because scoring every compound in these screening libraries with a co-folding model would take years (Figure 3), we controlled instead with a scrambled-target null, each active scored against a random incorrect target, which preserves any ligand-only signal while removing the correct protein match, and reserved direct comparison to co-folding for the focused libraries where computation is tractable (Figure 4E, Table 2). Recognition of even the most novel ligands stayed far above both random and this null, which by adjusted logAUC sits at essentially random across the whole novelty range (about 0.01). The persistent gap between the Ptarmigan-1 curve and this null supports target-specific recognition, and it extends into the least-precedented regime. Restricted to the held-out targets (as in panel A), examples simultaneously novel in both chemistry and protein space retained a positive mean adjusted logAUC of 0.13 (*n* = 6; Figure 4C).

**Figure 4:**
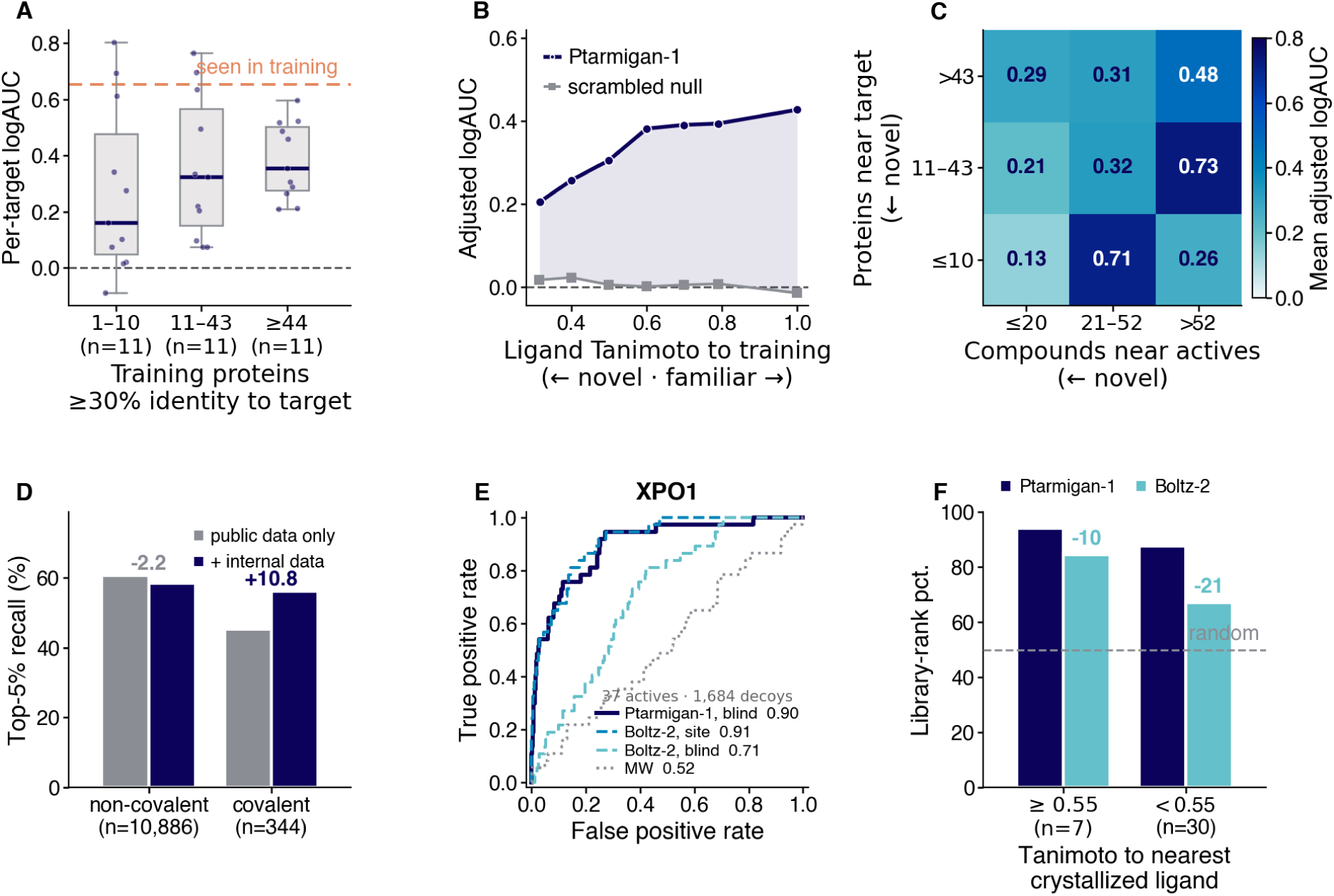
Screening performance as a function of target novelty and ligand novelty. “Held out” denotes a target whose protein sequence never appears in training; adjusted logAUC is scaled so a random ranker scores 0 and a perfect ranker approaches 0.855, and “top 5%” (panel D) is the fraction of actives ranked among the top 5% of the screened library. (A) Per-target adjusted logAUC for the 33 held-out targets of the screening benchmark (Supplementary Figure 4), binned into thirds by training-protein coverage sharing at least 30% sequence identity with the target (*n* = 11 targets per bin; boxes give median and interquartile range, points are individual targets). The upper dashed line marks the median adjusted logAUC of targets seen in training (0.65, the same benchmark), the grey dashed line marks random (0). (B) Active compounds of the external test set, binned by ECFP4 Tanimoto to the nearest training compound. Pooled adjusted logAUC of the actives in each bin for Ptarmigan-1 (navy) and for a scrambled-target null (grey). (C) Mean adjusted logAUC over a 3×3 grid of protein novelty (rows: training proteins within 30% identity of the target) and ligand novelty (columns: training compounds within an ECFP4 Tanimoto of 0.4 of the target’s actives), each binned into thirds over the 33 held-out benchmark targets (as in panel A); cell counts range from 2 to 6 targets. Color encodes the mean adjusted logAUC (D) Top-5% recall for held-out actives under a model trained on public data alone (grey) versus with the internal corpus added (navy), split by non-covalent (*n* = 10,886) and covalent (*n* = 344) actives. (E) Receiver-operating-characteristic curves for 37 covalent SINE actives against 1,684 property-matched decoys, scored by Ptarmigan-1 from sequence with no pocket (AUROC 0.90), Boltz-2 given the binding site (0.91), Boltz-2 blind to the binding site (0.71), and a molecular-weight baseline (0.52). (F) Library-rank percentile of the same 37 actives, split by ECFP4 Tanimoto to the nearest SINE co-crystallized with XPO1 in the PDB (*n* = 7 at ≥ 0.55, *n* = 30 at < 0.55).

**Table 2:**
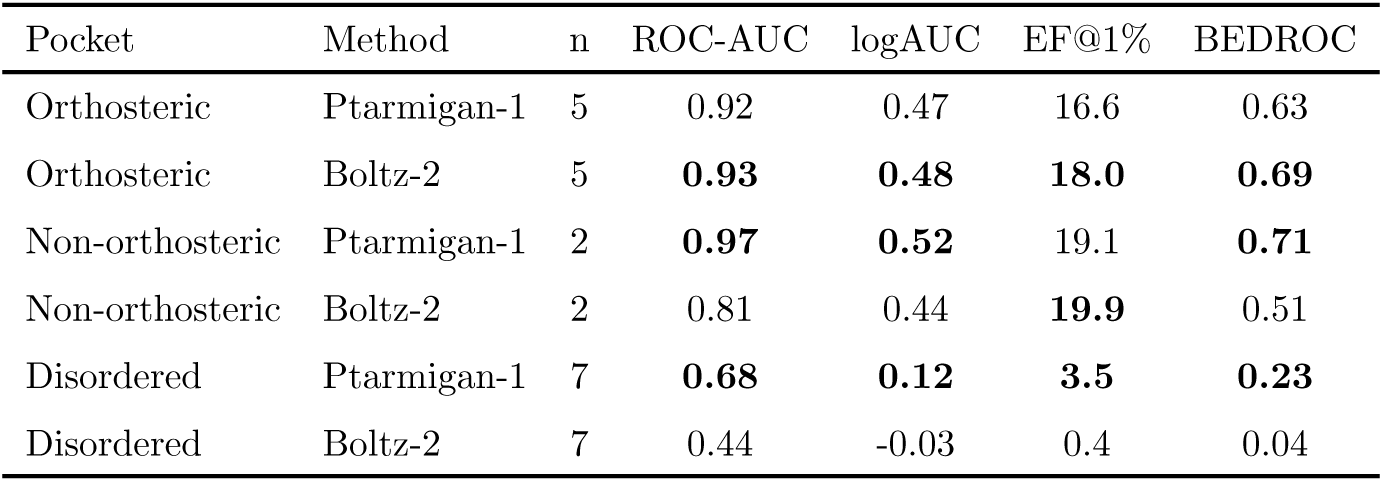
Covalent virtual-screening performance by pocket class. For each pocket class (the organizing axis of Supplementary Figure 5) and method, entries are means over the *n* targets in that class: ROC-AUC, adjusted logAUC (*λ* = 0.001, random = 0), enrichment factor at 1% (EF@1%), and BEDROC (*α* = 20); within each pocket class the higher value in each metric is set in bold. The classes are orthosteric (*n* = 5), non-orthosteric (*n* = 2; KRAS and XPO1), and disordered (*n* = 7); per-target values with bootstrap 95% confidence intervals are given in Supplementary Figure 5B. Within each target both methods scored an identical set of compounds, every active retained, with each score oriented so that higher indicates greater predicted activity (Methods).

Given that Ptarmigan-1’s covalent reach draws on residue-level chemoproteomic data (Supplementary Figure 1), we asked how much of its generalization depends on our internal corpus by comparing the full model against one retrained on public data alone. Top-5% recall for non-covalent actives was essentially unchanged (a 2.2-point decrease with the internal corpus added), but for covalent actives the internal corpus lifted recall by 10.8 points, from 45% (public-only) to 56% (Figure 4D), suggesting that the internal covalent data slice helps extend generalization to covalent chemistry.

Finally, we examined one held-out target in detail, XPO1, whose covalent inhibitors engage a non-orthosteric cysteine rather than an orthosteric pocket. Against 37 selective inhibitors of nuclear export (SINEs) and 1,684 property-matched decoys, Ptarmigan-1 scored compounds from sequence alone and reached an AUROC of 0.90. Boltz-2 weakly scored XPO1 actives when the binding site was not pre-defined (0.71), and rose to comparable performance with a fixed, predefined covalent binding site (0.91; Figure 4E). The two models generalize differently across chemical space: Boltz-2 ranked the SINEs most similar to those already co-crystallized with XPO1 in the PDB well above the more novel analogs (library-rank percentile 85 versus 67), whereas Ptarmigan-1 ranked both near the top of the library (94 versus 88), recovering novel chemistry that a structure-based method recognizes mainly where a solved complex already exists (Figure 4F).

### Ptarmigan-1 retains enrichment on non-orthosteric and pocketless targets

The targets that most motivate structure-free screening are those without an annotated canonical pocket, but benchmarks for this task remain underdeveloped. To assess Ptarmigan-1 here, we built a covalent benchmark combining ChEMBL and internal covalent binding data, held out from training. We assessed Ptarmigan-1 against Boltz-2, a representative open-weight co-folding model, on this benchmark. On targets with a canonical orthosteric pocket, the two methods were comparable (Table 2). On the two targets categorized as non-orthosteric pockets, Ptarmigan-1 showed stronger average performance, though this category holds only two targets, on which the two models diverge sharply. As described above, Ptarmigan-1 recovered and ranked XPO1 SINEs better. On the KRAS switch-II pocket, a non-orthosteric site well-characterized in the structural literature, Boltz-2 outperformed Ptarmigan-1 (Supplementary Figure 5). On the intrinsically disordered targets (AlphaFold2 pLDDT < 50), Boltz-2 fell to near-random while Ptarmigan-1 retained positive enrichment across the targets.

### Ptarmigan-1 identifies cryptic pockets

For traditionally “undruggable” proteins, identifying a cryptic, ligandable binding site is often a necessary and challenging first step in drug discovery [47, 48]. We hypothesized that we could leverage the generalization and speed provided by Ptarmigan-1 to identify cryptic sites without explicitly training the model for this task. To test this hypothesis, we screened a diverse compound library against proteins with known cryptic sites, scored every residue of these targets, and used the strongest predicted engagement across the protein to identify cryptic ligandable sites. Unlike other reported approaches for this task, this would act as a blind pocket predictor that needs neither a resolved 3D structure nor a bound ligand. We evaluated this on CryptoBench [49], a benchmark of cryptic binding sites absent from the unbound (apo) structure that open only upon ligand binding. We used its held-out test split of 222 apo structures (199 scored) and evaluated per-protein AUPRC.

Scoring zero-shot from sequence, Ptarmigan-1 recovered the cryptic-site residues better than P2Rank [45], the structure-based pocket finder given the apo structure (per-protein AUPRC 0.27 versus 0.22), which we ran ourselves on the same 199 structures under the same metric (Figure 5A); Ptarmigan-1 was never trained to predict pockets. For context, [49] reports 0.36 for its own supervised network, trained on this benchmark’s cryptic sites from ESM2-3B embeddings, and 0.19 for PocketMiner [50]. Both are pooled (micro) per-residue AUPRC over that study’s own evaluable subsets rather than the per-protein values used here, so neither is a head-to-head comparison and we plot only the methods we scored identically. The advantage over the structure-based pocket finders is specific to cryptic sites. On the well-folded orthosteric pockets of COACH420 [45], the structure-based P2Rank led (0.63 versus 0.58; Figure 5B). Cryptic site recovery was independent of prior training exposure, with Ptarmigan-1 scoring nearly the same on test proteins never appearing in its training as on those it had seen (held-out 0.26 versus seen 0.28; Figure 5C). As an example, we show the apo structure of carboxypeptidase A, where predicted engagement concentrates on the residues lining the cryptic site even though no ligand is bound (Figure 5D).

**Figure 5:**
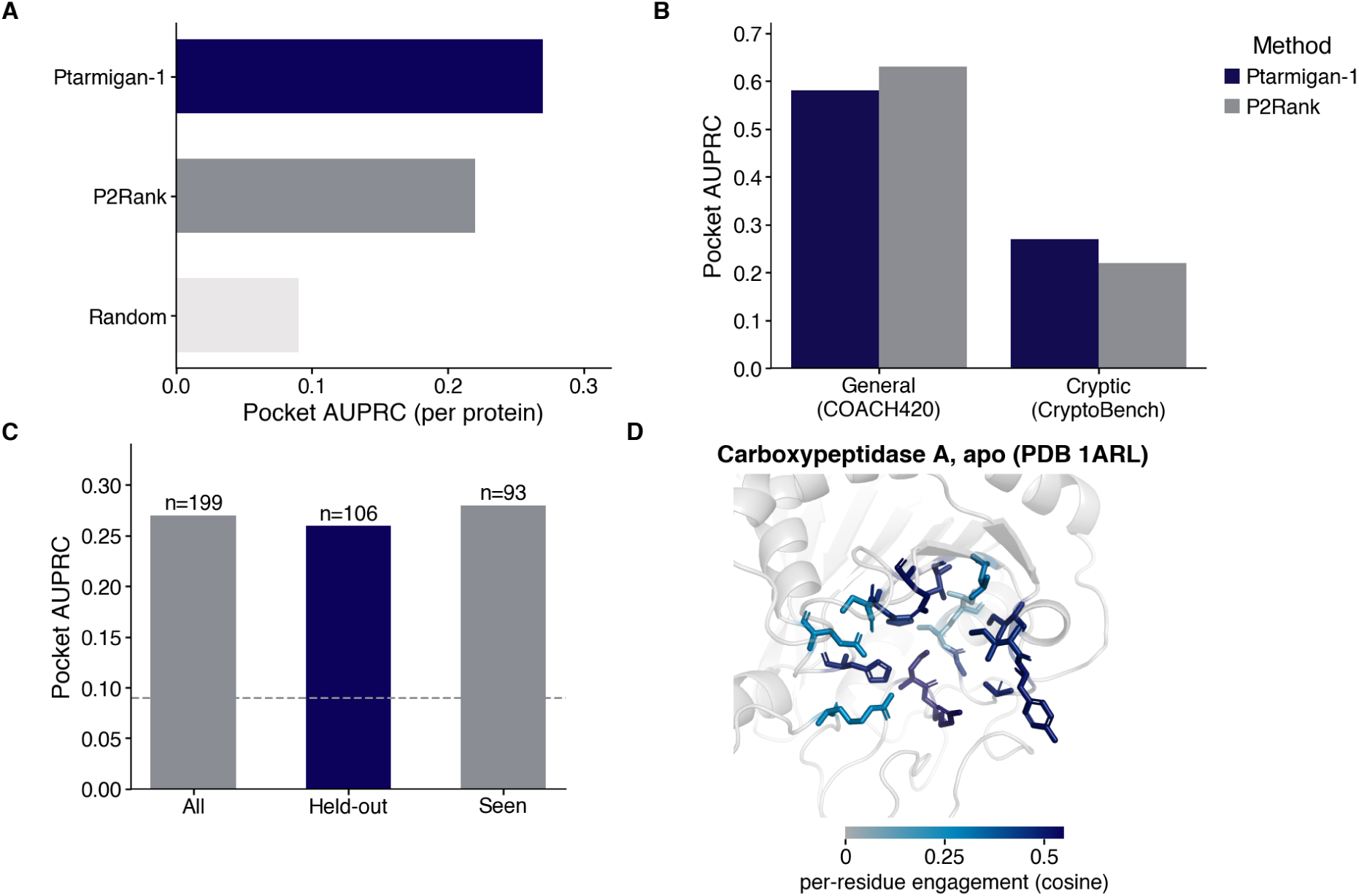
Blind cryptic-pocket identification from sequence. Per-residue engagement is marginalized over a compound library and scored as per-protein AUPRC over residues ranked by ligandability (positives are residues within the annotated site). (A) Cryptic-site comparison on the CryptoBench test split (222 apo structures, 199 scored). Every bar is computed here on the same 199 proteins under per-protein AUPRC: Ptarmigan-1 (indigo) scoring zero-shot from sequence, P2Rank (grey) given the apo structure, and a random ranker. The values [49] reports for its own supervised pLM-NN (0.36) and for PocketMiner (0.19) use a pooled (micro) per-residue AUPRC over that study’s evaluable subsets; they are given as context in the main text rather than plotted here, since that metric and protein set would not be comparable with these bars. (B) Per-protein AUPRC for Ptarmigan-1 and P2Rank on two benchmarks: the well-folded orthosteric pockets of COACH420 and the cryptic sites of the CryptoBench test split, where the pocket is absent from the apo structure. Both methods use the same metric on the same structures within each benchmark. (C) Ptarmigan-1’s per-protein AUPRC split by whether the test protein’s UniProt accession appears in training (*n* = 106 absent, *n* = 93 present); the dashed line marks a random ranker. (D) Worked example: the apo structure of carboxypeptidase A (PDB 1ARL), with the annotated cryptic-site residues drawn as sticks and colored by Ptarmigan-1’s per-residue engagement (grey low to indigo high) over a translucent cartoon. No ligand is present in the structure.

### Ptarmigan-1 recovers recent STAT6 inhibitors and localizes them to the SH2 domain

To test model performance on a target and compound set outside of its training data, we asked whether Ptarmigan-1 could recover binders of the STAT6 transcription factor from a set of patent-disclosed actives. We selected two disclosed inhibitor series, a Pfizer series across two patents [51, 52], whose binding site is not disclosed, and a Recludix series disclosed to occupy the orthosteric SH2 phosphotyrosine pocket [53] (Figure 6). 40 diversity-picked exemplars were selected from each each. All three patents were published after Boltz-2’s structural training cutoff [9], and a compound-by-compound check against every source in our own training corpus found no exact structural match to any of the 80 actives (median nearest-neighbor ECFP4 Tanimoto 0.35 Pfizer, 0.31 Recludix; Figure 6A), and no structurally similar compounds are linked to STAT6 in our training data. No exact training-set match therefore explains either model’s behavior on these series (Supplementary Figure 6), though neither this nor the temporal cutoff rules out learned chemotype, target-family, or assay-source correlations.

**Figure 6:**
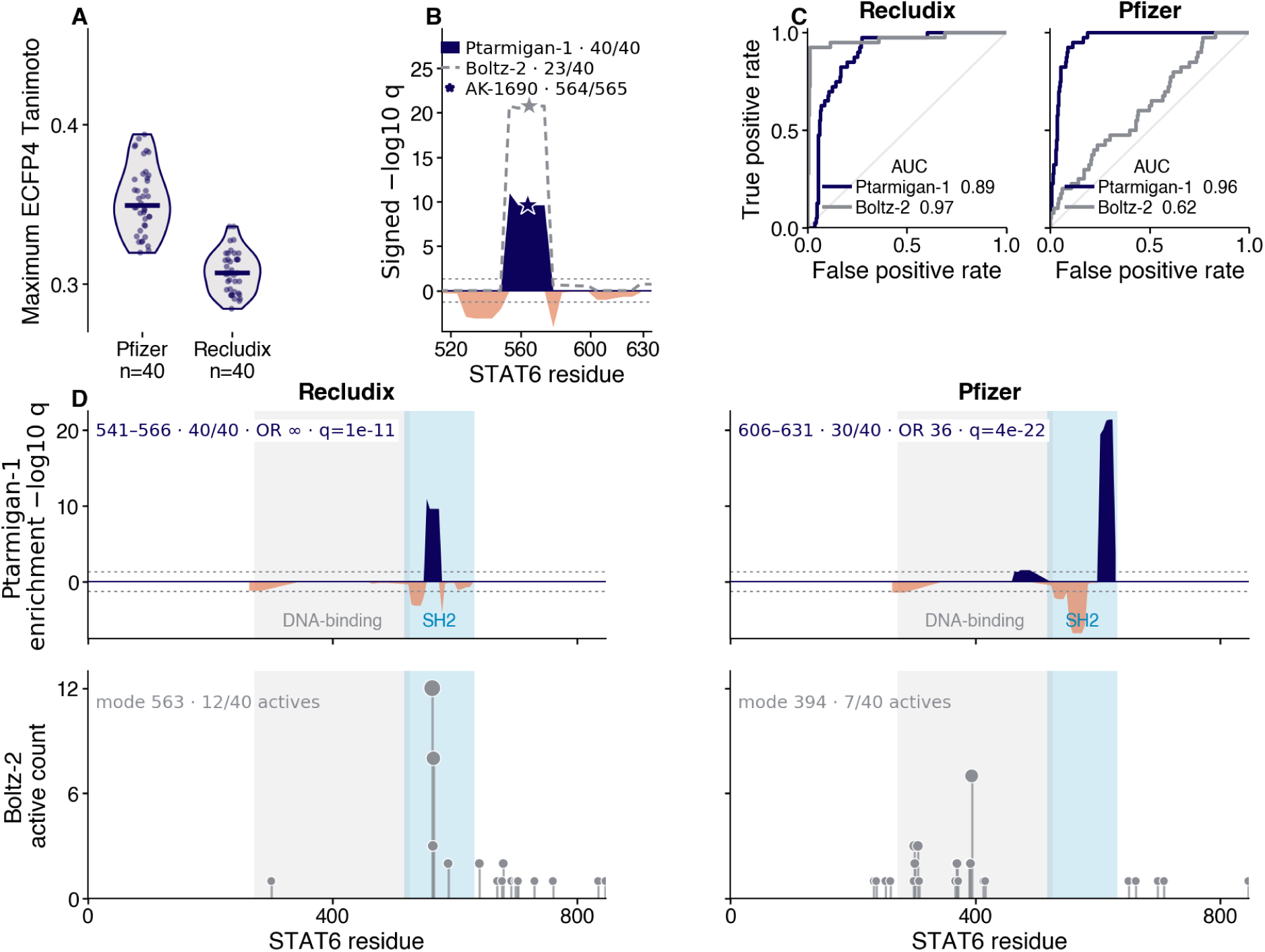
Recovery and predicted localization of patent-disclosed STAT6 inhibitors. (A) Maximum ECFP4 Tanimoto of each active to the nearest training compound (violin and points; bar, median). (B) Orthosteric validation on the AK-1690. Signed −log10 q of a moving-window enrichment test across the SH2 region (residues 515–635) for Ptarmigan-1 top-engagement residues (filled) and Boltz-2 predicted contacts (dashed), each against its decoy null. (C) Screening ROC for each series and model against property-matched decoys. (D) Predicted localization along the full STAT6 sequence (UniProt P42226, 847 aa). Top: Ptarmigan-1’s moving-window enrichment (signed −log10 q; positive where actives concentrate above decoys), with the strongest FDR-significant window annotated (Recludix 541–566, 40/40 actives, odds ratio ∞, *q* = 10^−11^; Pfizer 606–631, 30/40, odds ratio 36, *q* = 4 × 10^−22^). Bottom: count of Boltz-2 predicted top contact residues for the 40 actives (mode 563 Recludix, 394 Pfizer). Shaded bands, the DNA-binding (273–526) and SH2 (517–632) domains.

On the Recludix series, whose orthosteric SH2 site is disclosed, both models separated actives from property-matched decoys (Ptarmigan-1 AUC 0.89, Boltz-2 0.97) (Figure 6B, Figure 6C). On the Pfizer series, whose site is undisclosed, Ptarmigan-1 maintained strong recovery of known actives enrichment (AUC 0.96) while Boltz-2 fell near random (0.62) (Figure 6C).

We then considered site localization across the two series. The established small-molecule site on STAT6 is its SH2 domain, a shallow phosphotyrosine-binding pocket through which STAT6 is recruited to activated IL-4 and IL-13 receptors and homodimerizes. Cell-permeable phosphopeptide-mimetic inhibitors that occupy this pocket block Tyr641 phosphorylation and downstream transcription [54], and the only publicly available cocrystal of a small molecule bound to STAT6 shows AK-1690, a phosphotyrosine-mimetic ligand in this pocket [55] (a structure deposited after the Boltz-2 training cutoff) As a positive control, we localized AK-1690 itself, which is also structurally novel to the Ptarmigan-1 training set (nearest-neighbor ECFP4 Tanimoto 0.36) yet has an experimentally resolved binding site. Ptarmigan-1 placed its peak engagement at residue 564, squarely within the SH2 phosphotyrosine pocket, recovering the known site of the compound (Figure 6B).

Ptarmigan-1 localized the Recludix actives to the phosphotyrosine pocket (window 541– 566, 40/40 actives, *q* = 10^−11^), coincident with the AK-1690 control and consistent with their disclosed orthosteric mechanism (Figure 6B, Figure 6D). Poses modeled by Boltz-2’s fell correctly within the SH2 pocket as well (mode residue 563).

For the Pfizer actives, Ptarmigan-1 localized the series instead to a distinct C-terminal SH2 sub-region (residues 606–631, 30/40 actives, *q* = 4 × 10^−22^; Figure 6D). Boltz-2 by contrast placed its predicted contacts in a separate region centered in the coiled-coil and DNA-binding region (mode residue 394; Figure 6D), along with several other putative pockets across the Pfizer series. These divergent predictions provide testable hypotheses for future biochemical or crystalization experiments on STAT6 and its inhibitors.

## Discussion

### Faster screening for traditional, structured targets

Three-dimensional structure is an intuitive artifact for human interpretation of compound– protein interactions, but our results suggest that computational models for small-molecule discovery do not require an explicit structure for predictive accuracy or generalization. Building those structures is also computationally expensive, limiting the practical scale of virtual screening. Ptarmigan-1 engagement reduces screening to a single matrix multiplication, allowing the model to economically screen billions of compounds against the whole proteome and resolve each prediction to the residues a compound is likely to engage.

That speed is valuable, and largely complementary to structure-based methods on well-folded targets. No benchmark offers a perfect comparison between the two architectures, but Ptarmigan-1 ranks actives comparably to a collection of structure-based methods on well-structured pockets (Table 1). The trade-off for these structure-free models is that they report no explicit binding pose, instead nominating the likely pocket that a compound engages.

We therefore see structure-free and structure-based screening as complementary. Ptarmigan-1 can narrow billions, eventually trillions, of compounds for a single target, and counter-screen them across the whole proteome to bias for selectivity and polypharmacology. Then that prioritized, tractable sub-library can be scored by a high-fidelity structural engine that resolves poses in physical detail and ranks compound affinities. This hybrid is already foreshadowed by methods that graft affinity heads onto docked poses [56], and grows more compelling as co-folding accuracy advances [24]. The broader and cheaper the upstream filter, the more a slow, accurate structural refiner can afford to spend on each surviving compound.

### Unlocking novel pockets and chemotypes for challenging targets

Ptarmigan-1’s distinctive capability lies at the sites where an explicit pose is hardest to build. On the covalent benchmark (Table 2) it was comparable to Boltz-2 on non-orthosteric cryptic sites and pulled clearly ahead on intrinsically disordered proteins.

Speed helps with disordered pocket screening as well. By screening multi-billion-compound libraries, Ptarmigan-1 shows an emergent ability to identify cryptic sites more efficiently than established methods, without ever being explicitly trained for that task (Figure 5). From sequence alone it localized two recent inhibitor series to the shallow SH2 site of STAT6 (Figure 6), a site a co-folding model recovered only for the series whose orthosteric pose it could resolve.

Ptarmigan-1 learns this breadth from multi-modal training data (reactive-cysteine chemoproteomics and activity-based profiling alongside conventional bioactivity, at mixed resolution) that structure-based models cannot readily use. As different assays expose different modes of engagement, we expect the model to learn a wider range of pocket behavior than the orthosteric, well-folded distribution the Protein Data Bank imposes on structure-based methods, and we expect it to bring covalent sites, allosteric grooves, and disordered regions within reach of virtual screening, ultimately discovering new pockets for historically “undruggable” targets.

### Moving beyond screening to compound look-up

In the near term, we are pursuing prospective experimental validation and a public web server for Ptarmigan-1. Beyond that, we believe a structure-free approach changes what a virtual screen fundamentally is. Docking and co-folding score every target and library from scratch, so each screen is a fresh computation whose reach is fixed in advance by the chemistry one decides to enumerate and the compute one can afford. When engagement is instead a distance in a learned embedding, that embedding is computed once and reused. Each new target or library becomes a lookup rather than another full screen, and the space itself improves as data accumulate, since every new measurement refines the space and sharpens future queries.

In that paradigm, compute and the choice of chemical space stop being the key constraints. What limits discovery becomes how quickly the compounds a model nominates can be synthesized, assayed, optimized, and advanced forward toward eventual preclinical and clinical evaluation.

## Methods

### The Ptarmigan-1 model architecture

Ptarmigan-1 co-embeds protein residues and small molecules into a shared vector space, so that a residue lies close to the compounds that engage it. We encode protein sequences with ESM-C, the 600M-parameter ESM Cambrian model [40], and compounds, from their SMILES string, with ChemBERTa [41]. Both are foundation models pretrained on sequence and chemical corpora alone, neither of which is biased toward well-folded proteins or structurally resolved ligands. We take the final-layer embeddings from ESM-C as one vector per residue, discarding the flanking start- and end-of-sequence tokens, and pool ChemBERTa at its classification token to a single compound vector. During training we cap protein inputs at 768 residues and tile longer sequences into overlapping windows, so that engagement anywhere along a long protein remains learnable. At inference a protein is encoded in a single forward pass under a length cap, taken from the checkpoint configuration where set and otherwise twice the training cap, and residues beyond that cap are not scored. Windows are therefore a training-time device and are not recombined at inference; the benchmarks reported here use a cap of 1,400 residues. A per-encoder linear projection maps both into a shared, *D* = 256-dimensional space, and we *L*_2_-normalize every embedding so that the similarity *s* between a residue and a compound is their dot product (cosine similarity). To preserve what the foundation models already encode, we keep both transformer backbones frozen and fine-tune only low-rank adapters (LoRA), a parameter-efficient fine-tuning method [57]; the adapters (rank 32, *α* = 64, dropout 0.1) are inserted into the attention and feed-forward projections of each encoder, parameterized separately for the two backbones, and trained together with the two projection heads.

We map the residue–compound similarity *s_r_* at residue *r* to a per-residue engagement probability 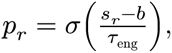 where *σ* is the logistic function and the bias *b* and temperature *τ*_eng_ are scalars learned jointly during training. A protein-level engagement probability follows from a temperature-scaled softmax pooling of the residue similarities, so that a small number of strongly engaged residues drive the protein-level call largely independently of sequence length. This aggregation gives Ptarmigan-1 flexibility in its training data: because a protein-level prediction requires no residue-level labels, we can train from sources that report only whether a compound engages a protein alongside those that resolve the engaged residue.

### Training Ptarmigan-1 to co-embed ligands with their target residues

We train Ptarmigan-1 by minimizing a weighted sum of complementary contrastive and calibration objectives, each chosen to exploit a different kind of label. A residue-level contrastive loss, in the style of CLIP [58], aligns each labeled binding residue with its paired compound against the other compounds in the minibatch. This term is available only for the sources that resolve binding to individual residues.

A protein-level contrastive loss applies a symmetric cross-entropy over the matrix of protein–compound engagement probabilities, treating the other compounds in the batch as in-batch negatives. Because this term needs only a binary engagement label it is where the bulk of our bioactivity data enters training. We pair this with an in-batch calibration loss, a binary cross-entropy in which the diagonal pairs are supervised by their measured engagement label and the off-diagonal pairs by zero, so that the protein-level probability reads as a calibrated value rather than a bare ranking score.

We optimized all trainable parameters with AdamW [59] (learning rate 6 × 10^−4^, weight decay 0.01) under a one-cycle learning-rate schedule [60] in bf16 mixed precision, for 2.5 × 10^4^ steps with gradients clipped to unit norm. The four objectives are combined with equal weight (residue-level contrastive, protein-level contrastive, residue-pair, and calibration terms all at 1.0); the covalent term is disabled for this model. Minibatches hold 256 protein–compound pairs per device across four devices under distributed data parallelism, giving an effective batch of 1,024, and in-batch negatives are gathered across devices so every pair is contrasted against all 1,023 others rather than only the 255 on its own device. Residue-resolved rows are oversampled by a factor of 50 relative to their frequency, since they are a small minority of the corpus but supervise the residue-level objective.

Validation is drawn from the training rows rather than from a separate file, by holding out 2% of Bemis–Murcko scaffolds and, independently, 2% of protein identifiers, and is evaluated every 500 steps. No benchmark row is used for validation. Training runs the full step schedule without early stopping, and the model reported throughout is the final checkpoint of that schedule rather than one selected on the validation metric or on any downstream benchmark.

### Training data sources

Each training example is a (protein sequence, compound SMILES) pair carrying per-residue labels (engaged, non-binding, or unknown) together with a protein-level engagement label and, where measured, covalent annotations. To measure generalization to new chemistry, we partition compounds by their Bemis–Murcko scaffold [61] and hold out 5% of scaffolds as a test set, so that no scaffold is shared between training and test. We further guarantee that every evaluation benchmark is unseen during training by routing a row to the test set whenever its compound matches a benchmark compound by InChIKey skeleton, which also catches protomers, tautomers, and stereoisomers, or its protein matches a benchmark target by UniProt accession [62].

We harvested protein–ligand complexes from the Protein Data Bank [16], retaining drug-like ligands while discarding cofactors, crystallization additives, and polymeric ligands. For each complex we labeled the residues within 5 Å of the ligand as engaged.

We drew protein-level engagement labels from public bioactivity resources: BindingDB [18]; the integrated kinase-inhibitor bioactivity set KIBA [63]; and LCIdb, a large consensus compound–target interaction database [64]. For each, we labeled a compound as engaging its target below an affinity threshold of 10 µM and as non-binding otherwise, without residue-level labels.

External covalent, residue-resolved labels came from four competitive activity-based protein profiling (ABPP) studies [30, 31, 65, 66]. For the studies reporting proteome-wide competition ratios, we labeled cysteines with high competition as engaged and those with low competition as non-binding, yielding genuine residue-level negatives; the remaining studies contribute engaged sites only.

We augmented these public sources with an internal corpus from our regulome profiling platform, a chemoproteomic assay that measures target engagement directly in native samples. These measurements supply additional residue-level engagement labels and comprise the bulk of residue-level labels in our dataset.

### Efficient retrieval with approximate nearest-neighbor indexing

Because Ptarmigan-1 scores a protein–compound pair by the dot product of *L*_2_-normalized embeddings, virtual screening reduces to nearest-neighbor retrieval. We embed a compound library once and store the vectors in a vector database (LanceDB) under an inverted-file product-quantization (IVF-PQ) index [67], which trades exact distances for sublinear query time. To screen a target, we encode its residues and query the library for the compounds nearest each residue, scoring every candidate by its per-residue similarity and the pooled protein-level engagement. The same index, built instead over an embedded proteome, lets us invert the query and rank proteins against a single compound to estimate proteome-wide selectivity.

### Proteome-wide screen and its matched null

Each per-protein score is a maximum over billions of compounds; thus, a high value is expected by extreme value alone, independent of any learned engagement. To separate genuine engagement from this library-size effect, we recomputed the screen under a matched null that holds everything fixed except the learned cross-modal alignment. We drew a fixed Haar-uniform orthogonal matrix *R* ∈ *O*(256) from a single seed and applied it to every residue embedding, *r* → *Rr*, while leaving the compound library untouched, then repeated the identical maximum-over-library selection. Because *R* is orthogonal it preserves each residue’s norm and every residue–residue inner product, so the geometry of the protein embedding cloud, and with it all of the sequence information ESM-C encodes, is retained exactly; only the orientation of the residues relative to the compounds, the alignment the contrastive objective learned, is destroyed. The rotated screen is therefore an extreme-value baseline for the same library under the same selection with the learned alignment removed, so the gap between the real and rotation-null best-compound distributions (Figure 1D) isolates the contribution of that alignment. A single rotation gives each protein only one null realization, which supports comparing the two distributions but not testing an individual protein. We therefore repeated the null on a stratified subsample of 1,000 proteins under 40 independently drawn rotations, sampled across bins of mean pLDDT, sequence length, and protein family, and deliberately over-representing proteins with a mean pLDDT below 50 (177 of the 1,000) so that the least structured proteins carry their own statistics. Because rotating a residue embedding by *R* and rotating a compound embedding by *R^T^* give the same similarity, all 40 rotations are evaluated within a single pass over the library rather than one pass each. For each protein we report the empirical tail probability (1 + *k*)/(1 + 40), where *k* is the number of rotations whose best-compound score reaches the protein’s real score, and control the false discovery rate across the subsample with the Benjamini–Hochberg procedure. Each rotation applies one global *R* to every protein, so null draws are correlated across proteins within a rotation.

### Evaluation datasets

To evaluate Ptarmigan-1 on conventional, non-covalent virtual screening, we used LIT-PCBA, a benchmark of dose-response PubChem assays curated to remove assay artifacts and to match the physicochemical properties of its actives and inactives [68]. We report the five target-sets for which published structure-based baselines are available, the estrogen-receptor-α agonist and antagonist sets, MAPK1, PPARG, and TP53, and compare against the per-compound Boltz-2 [9], Protenix, Glide-SP, Gnina-Vina, and Gnina-CNN scores released by Shen et al. [42]. We additionally ran two co-embedding baselines ourselves on the same compounds: ConPLex [33], which scores from protein sequence and a molecular fingerprint using its released BindingDB checkpoint, and SPRINT [34], which co-embeds a molecular fingerprint with a SaProt structure-token encoding of the target that we derived from each protein’s AlphaFold model, using the authors’ LIT-PCBA checkpoint. We scored both with the same metric battery as Ptarmigan-1, so their numbers are directly comparable; we excluded DrugCLIP [32], which requires a three-dimensional binding pocket and reports enrichment only through its own pipeline, without a comparable adjusted logAUC (Supplementary Table 2). Every method scored an identical set of compounds, each score oriented so that higher values indicate greater predicted activity, so that differences reflect ranking rather than coverage.

Across the covalent and disordered benchmarks that follow, Boltz-2 is our sole structure-based comparator. It is the only co-folding model that both scales to a full-library screen and ranks compounds directly through an affinity head, rather than emitting only a predicted structure and confidence as AlphaFold3, Protenix, and Chai-1 do; co-folding a pose for every library compound is otherwise prohibitive at this scale [42]. The degradation of structure-based methods on non-orthosteric and disordered sites is architecture-independent across AlphaFold3, Protenix, Boltz-2, Chai-1, and DynamicBind [21], so Boltz-2 stands in for the class; newer systems reporting better generalization are closed and untested at library scale, so they cannot serve as a screening baseline here [24].

To assess residue-level localization, we used COValid, a set of covalent inhibitors with experimentally mapped engaged cysteines [43]. It comprises 874 covalent actives spanning ten cysteine sites across nine proteins, each active annotated with the single cysteine it modifies; we scored every active, ranked all residues of the target by engagement, and took the percentile of the annotated cysteine as the readout.

To extend the localization test to non-covalent ligands, we used the PoseBusters benchmark set (version 1), 428 recent protein–ligand complexes assembled to postdate the training cutoffs of common docking and co-folding methods [44]. We mapped each complex to a UniProt accession through the RCSB and labeled it seen or held out by whether that accession appears in our training splits. Because the set mixes cofactors, oligosaccharides, and peptide-like ligands with drug-like small molecules, we restricted quantitative comparisons to a drug-like subset defined by an RDKit filter on molecular weight, heavy-atom count, and sugar content, leaving 337 complexes, of which 330 carry a training-seen label (288 seen, 42 held out). For each complex we scored every residue from sequence alone and defined the pocket as the residues within 5 Å of the crystallized ligand. The four covalent worked examples (Figure 2A) were scored the same way, with pockets taken from their co-crystal structures (BTK, PDB 5P9J; BMX, 8X2A; ITK, 9NWX; the KRAS G12C mutant, 6OIM); the structure insets paint PDB 5P9J (BTK, Figure 2B) and 7AS1 (influenza PB2 with pimodivir, Figure 2E) by engagement.

To test whether localization generalizes to recent, unseen systems, we used Runs N’ Poses [23], a benchmark of 2,815 protein–ligand systems deposited after the 30 September 2021 PDB cutoff and stratified by structural similarity to that cutoff. Because Ptarmigan-1 trains on binding assays rather than PDB structures alone, that temporal cutoff does not bound its training: 96% of these proteins fall within 30% sequence identity of a training protein and 98% of the ligands within an ECFP4 Tanimoto of 0.4 of a training compound. We therefore re-binned the systems by similarity to Ptarmigan-1’s own training set (sequence identity to the nearest training protein; ECFP4 Tanimoto to the nearest training ligand) and, for each system, scored every residue against the crystallized ligand and read out both top-residue recovery, whether the single highest-engagement residue lies within 5 Å of the ligand, and the pocket-vs-rest AUROC (Supplementary Figure 3).

For covalent screening we curated a benchmark from ChEMBL [17] spanning seven well-characterized covalent targets across two tiers: orthosteric cysteines at canonical or catalytic sites, namely EGFR (C797), ERBB2 (C805), FGFR4 (C552), cathepsin L (C25), and BRD4 (C356), whose covalent actives engage the acetyl-lysine pocket; and non-orthosteric cysteines away from the primary site, namely the KRAS G12C mutant (C12) and XPO1 (C528). We assembled this set for greater target diversity than the kinase-dominated COValid benchmark [43], taking the high-confidence covalent binders of each target as actives and restricting them to compounds first reported after the Boltz-2 training cutoff to limit leakage. We paired each active with 50 decoys, drawn in a 70:30 mix of two strategies. Property-matched decoys are ChEMBL compounds that fall within fixed windows of at least one active on molecular weight (±50 Da), calculated logP (±1), and the counts of hydrogen-bond donors, hydrogen-bond acceptors, and rotatable bonds (±2 each), so that enrichment cannot be explained by gross physicochemical differences alone. Warhead-matched decoys instead share a target active’s reactive warhead class but are measured binders of a different protein, a hard negative for whether the model recognizes target engagement rather than warhead reactivity. We drew every decoy from compounds never assayed against the target, since a measured non-binder is more useful as a training negative than as a presumed-inactive decoy, and required each decoy to differ in its Bemis–Murcko scaffold from all of the target’s actives.

XPO1′s case study (Figure 4E, Figure 4F) draws on a separate, larger benchmark assembled specifically for that comparison, 37 selective inhibitors of nuclear export (SINEs) against 1,684 property-matched decoys, rather than the ChEMBL covalent set above, where XPO1 carries only 4 annotated actives (Supplementary Figure 5B), too few on their own for a stable ROC curve.

For the disordered class, we curated seven intrinsically disordered or poorly structured proteins from our internal regulome-profiling corpus and paired each covalent active with warhead-matched decoys.

Finally, to measure generalization, we assembled a larger held-out screening benchmark of 100 targets drawn from the test splits of our public training sources, sampling as many targets whose UniProt accession never appears in training as were available (33 unseen, 67 seen) and pairing each active with up to 50 cross-target decoys (Supplementary Figure 4).

To assess cryptic-pocket discovery, we used CryptoBench [49], a benchmark of sites absent from a protein’s unbound (apo) structure that open only upon ligand binding. We scored its official test split of 222 apo structures, keeping the 199 whose curated cryptic-site annotation mapped cleanly onto the deposited structure’s residue numbering. For each apo protein, we obtained a ligand-agnostic ligandability score by marginalizing Ptarmigan-1′s per-residue engagement over a 200,000 compound diverse subsample of the OnePot library. We ranked residues by this score and computed the per-protein AUPRC against the annotated cryptic-site residues. P2Rank was run by us on the same apo structures under this identical metric; PocketMiner and CryptoBench’s own supervised pLM-NN enter as the values [49] reports for this test split, computed under its pooled (micro), per-residue AUPRC rather than ours. We assessed the equivalent well-folded-pocket comparison on COACH420 [45], scoring Ptarmigan-1 and P2Rank identically.

As a case study, we asked whether Ptarmigan-1 recovers recently disclosed STAT6 inhibitors (Figure 6). We took two disclosed inhibitor series, 40 diversity-picked exemplars each: a Pfizer series (WO2025133961A1 [51], WO2025215579A1 [52]), whose binding site is not disclosed, and a Recludix series (WO2026015868A1 [53]), disclosed to occupy the orthosteric SH2 pocket. All postdate the Boltz-2 structural training cutoff, and none matches a training compound (median nearest-neighbor ECFP4 Tanimoto 0.35 and 0.31 respectively, no exact match; Figure 6A, Supplementary Figure 6). We ranked each series against its property-matched decoys with Ptarmigan-1 from sequence alone and Boltz-2 run blind, with no pocket specified. Both models were evaluated on the decoys their runs returned; for the Pfizer series this restricts both to the 597 of the 826 property-matched decoys for which Boltz-2′s co-folding run returned an affinity. The 229 decoys Boltz-2 failed to fold are not a chemically random subset (median molecular weight 551 versus 508 for those it scored, Mann–Whitney *p* ≈ 10^−15^), so restricting one arm alone would compare different decoy sets; imputing the 229 failures as the worst- or best-ranked decoys brackets Boltz-2′s Pfizer AUC between 0.45 and 0.73. The Recludix decoys folded essentially completely (395 of 400). For localization we annotated the SH2 domain as UniProt P42226 residues 517–632 and used AK-1690, the phosphotyrosine-mimetic ligand of the only public STAT6 cocrystal (PDB 9BIG [55]), as a positive control. To localize each series along the sequence, we compared, in a window of 25 residues stepped by 5, the number of actives whose top-engagement residue (Ptarmigan-1) or predicted contact residue (Boltz-2) fell in the window against the same count over the pooled decoys, by a Fisher exact test. Per-window p-values were Benjamini–Hochberg adjusted across the sequence, and each window’s −log10 q was signed positive where the active fraction exceeded the decoy fraction and negative otherwise; windows are called significant at *q* < 0.05 (Figure 6B, Figure 6D). Panel B restricts this scan to the SH2 region for the Recludix series, whose site is known; panel D runs it over the full sequence for both series, with the lower track counting, per residue, the contacts Boltz-2 predicts across the 40 actives.

### Timing measurements

Throughput comparisons use wall-clock time on a single NVIDIA H100, and we report the quantity each measurement actually captures rather than a per-compound latency. For Ptarmigan-1 we timed one prediction run over the 5,488 unique EGFR ligands of the covalent benchmark against the EGFR sequence, in bf16 with a batch size of 32. Total wall time was 55.5 s, of which 1.9 s encoded the protein once, 6.1 s encoded the 5,488 compounds, and 0.5 s scored the pairs from the cached embeddings; the remainder is model loading and parquet input and output. Dividing total wall time by ligand count gives 10.1 ms per ligand, so compound embedding is included in that figure and the one-time costs are amortized over the run. For Boltz-2 we timed a co-folding shard over the same target, which completed 50 ligands in 44.6 min, or 53.5 s per ligand on the same amortized basis. The instrumentation recorded one start and one completion timestamp for the shard rather than per ligand, so no within-shard variance is available and the 50-ligand sample is small; the ratio of the two figures is accordingly quoted to one significant figure as roughly 5,000-fold rather than as a precise multiple. Both figures measure throughput under the batching each method uses, not the latency of a single isolated compound, and neither includes the one-time cost of embedding a screening library that is then reused across targets. Library embedding and retrieval scaling (Figure 3A, Figure 3B) were timed separately on the same hardware, sweeping library size with the compound encoder in bf16 and building the inverted-file product-quantization index on the resulting vectors.

### Evaluation metrics

Because virtual screening rewards the early recognition of true binders, we quantified ranking quality with the logarithmic AUC (logAUC), the area under the receiver-operating-characteristic curve integrated over log_10_ of the false-positive rate from a lower bound of *λ* = 10^−3^ [69]:

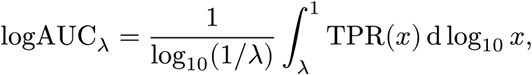

where *x* is the false-positive rate and TPR(*x*) the true-positive rate at that threshold. A random ranker satisfies TPR(*x*) = *x* and therefore scores (1 − *λ*)/(ln 10 ⋅ log_10_(1/*λ*)) ≈ 0.145 at *λ* = 10^−3^, while a perfect ranker scores 1. We report the adjusted form, which subtracts that random expectation:

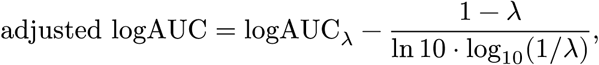

so a random ranker scores ≈ 0, a perfect ranker ≈ 0.855, and a ranker worse than random is negative. For virtual screening we computed this adjusted logAUC per target (ranking compounds against a protein); for localization we instead ranked a target’s residues by engagement and took the percentile of the annotated engaged cysteine among them.

For the non-covalent localization on PoseBusters we summarized each complex two ways. The pocket-vs-rest AUROC is the area under the ROC curve that ranks a protein’s residues by engagement with the pocket residues as positives, measuring how cleanly engagement separates the binding site from the rest of the chain. Binding-site recovery, the readout closest to the criterion structure-based methods report, records whether a complex’s single highest-engagement residue falls within 5 Å of the ligand; we report the fraction of the 337 drug-like complexes recovered, with an exact-binomial 95% confidence interval.

For the binding-site-recovery comparison (Figure 2G) we set Ptarmigan-1’s sequence-only recovery beside pose accuracy (pocket-aligned ligand RMSD < 2 Å) from the structure-based field. For AutoDock Vina [5] and DiffDock [46] we took the per-complex pose outcomes released with the PoseBusters study and recomputed the success rate on our drug-like subset, each method given the structure or pocket; for AlphaFold 3 [7] and Boltz-2 [9] we cite the published blind co-folding success rates on the full PoseBusters set.

To test whether localization depends on our internal corpus, we retrained Ptarmigan-1 on public data alone, using an identical architecture and training recipe with the internal chemoproteomic corpus removed, and compared its per-complex pocket-vs-rest AUROC against the full model’s on the held-out drug-like proteins (Figure 2H).

Pocket recovery alone cannot distinguish a compound-specific prediction from a protein-only ligandability prior, since both place the highest-scoring residues in the pocket most ligands occupy. To separate them we rescored the same 337 drug-like PoseBusters complexes under two controls and compared each against the correct pairing per complex (Supplementary Table 1). In the ligand-shuffle control, each protein is scored against a ligand belonging to a different protein, drawn to match the native ligand’s molecular weight (±50 Da), calculated logP (±1), and hydrogen-bond donor, acceptor, and rotatable-bond counts (±2 each), the same windows used for decoy construction, so that a loss in recovery cannot be attributed to a grossly different molecule; a donor is never taken from a complex sharing the query’s UniProt accession, and we average five independent draws within each complex before testing. In the ligand-agnostic control, per-residue engagement is marginalized over a fixed 24-compound probe panel, excluding each complex’s own ligand, giving the recovery available without any information about the query compound. For each readout we report the paired per-complex difference from the correct-ligand arm with a 95% percentile bootstrap interval over complexes and a Wilcoxon signed-rank test, since the absolute value of either arm alone is uninformative about conditioning.

## Data and code availability

The public training sources are BindingDB [18], KIBA [63], LCIdb [64], the Protein Data Bank [16], and four published activity-based protein profiling studies [30, 31, 65, 66]; each is available from its original publication or repository. The evaluation sets built on public data (LIT-PCBA, COValid, PoseBusters, Runs N’ Poses, CryptoBench, COACH420, and the ChEMBL covalent benchmark) derive from published resources, and we will release the compound identifiers, target accessions, labels, and decoy assignments used here, together with the per-figure source tables. The internal chemoproteomic corpus and the disordered-target benchmark drawn from it are proprietary and will not be released; every claim that depends on them is identified as such in the text, and the public-data ablations reported for localization (Figure 2H) and covalent recall (Figure 4D) quantify what changes without them.

## Supplementary Information

**Supplementary Figure 1:**
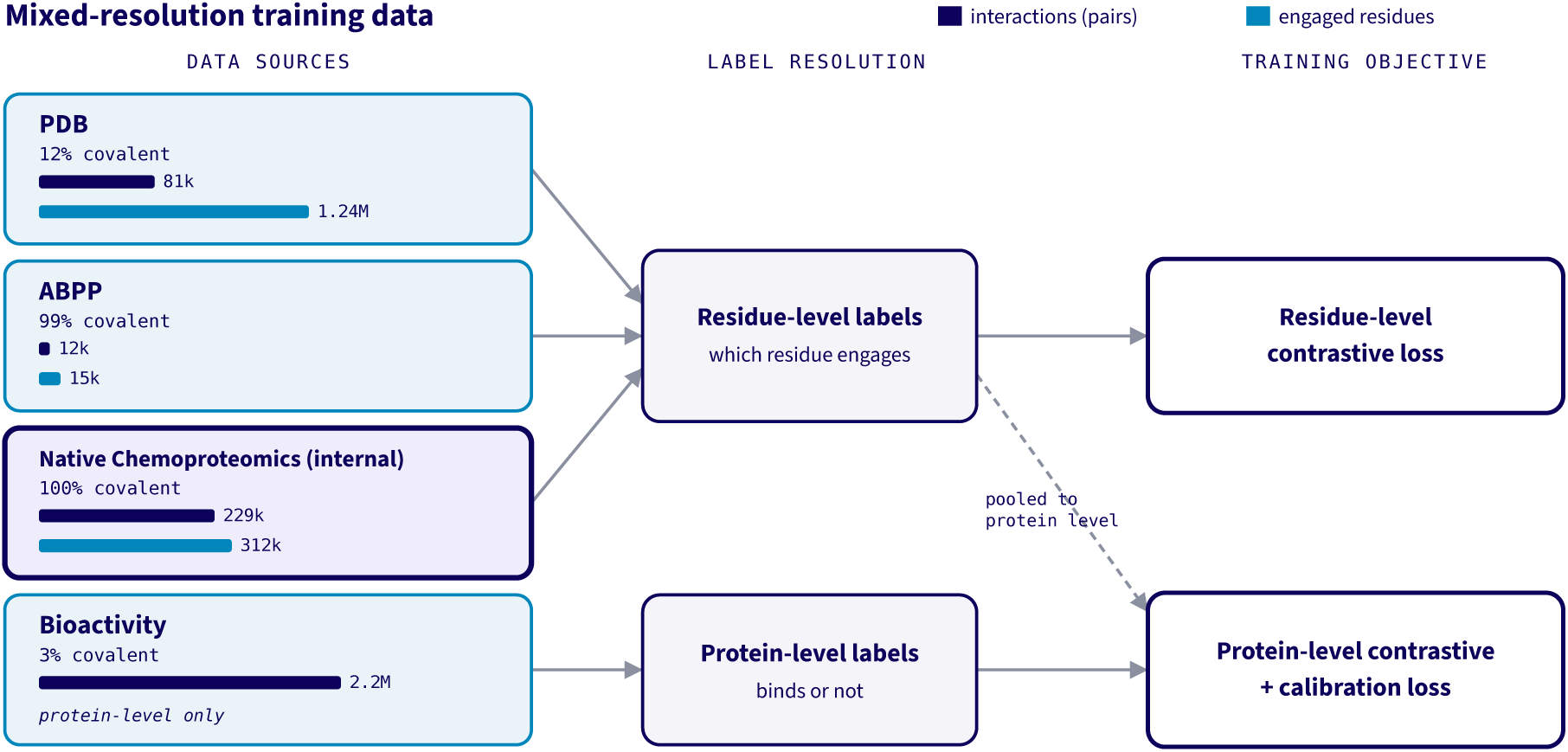
Ptarmigan-1 trains on protein–compound data of mixed resolution. Each source is drawn with the training objective it supervises, and bar length gives the number of protein–compound interactions on a logarithmic scale (2.6 million in total). Sources are grouped by label resolution. Protein-ligand complexes (PDB), reactive-cysteine chemoproteomics from published activity-based protein profiling studies (ABPP), and an internal native-chemoproteomic corpus resolve which residue a compound engages and so carry residue-level labels (blue bars give the count of engaged residues); the large bioactivity collections (BindingDB, KIBA, and LCIdb) record only whether a compound engages its target and are protein-level only. Residue-resolved sources train the residue-level contrastive objective and, pooled to the protein level, the protein-level objective, whereas bioactivity trains the protein-level objective alone. Border color marks provenance (public versus internal).

**Supplementary Figure 2:**
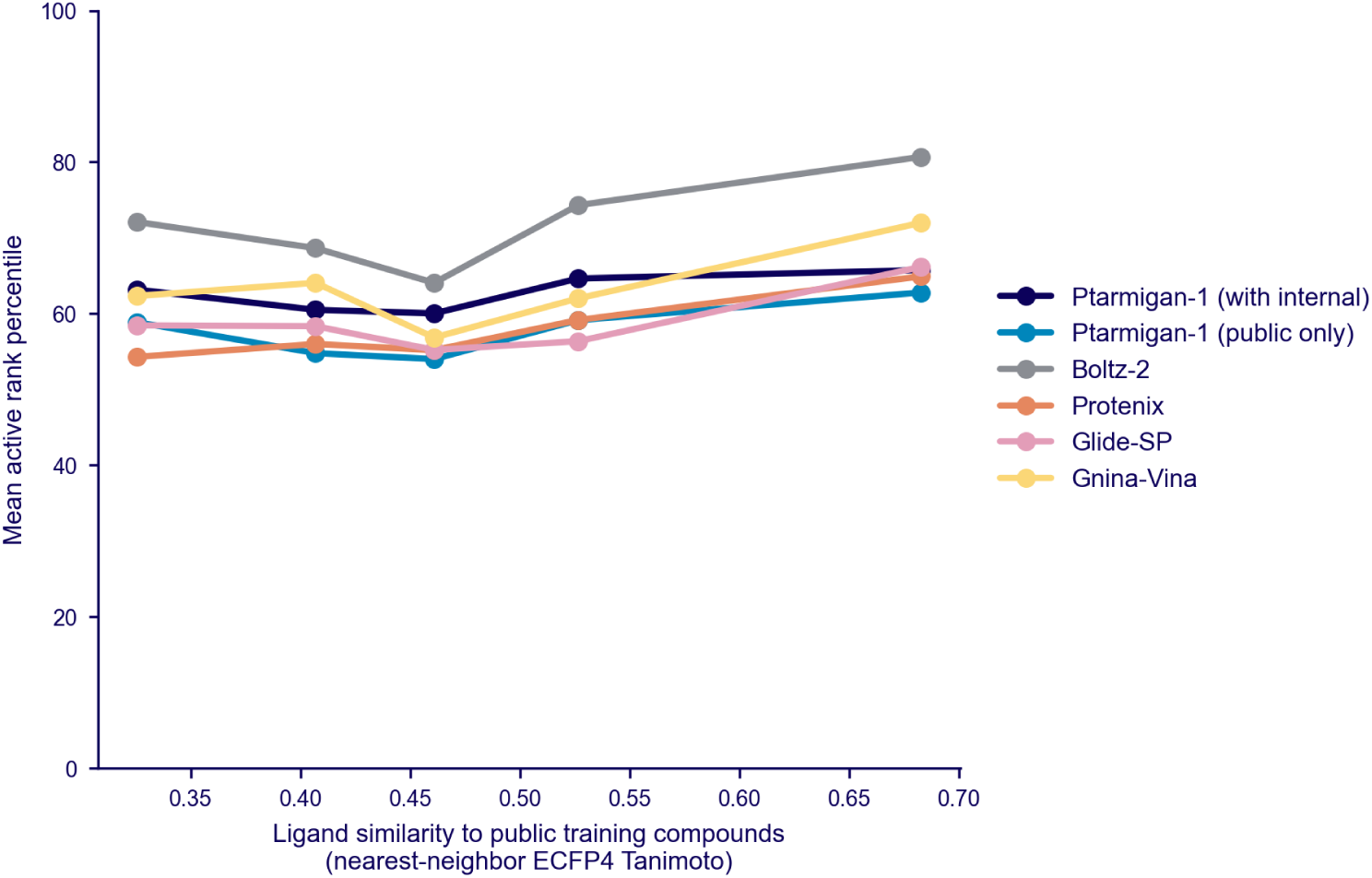
Recognition of LIT-PCBA actives as a function of ligand novelty, for Ptarmigan-1 and the published structure-based baselines. For each of the 529 actives across the five LIT-PCBA target-sets with published baselines (estrogen-receptor-α agonist and antagonist, MAPK1, PPARG, and TP53), we computed the nearest-neighbor Morgan (ECFP4) Tanimoto to the public training compounds (horizontal axis, binned into similarity quintiles) and, under each method, the active’s rank percentile among all compounds screened against its target. Each line is the mean active rank percentile per quintile, for Ptarmigan-1 with and without internal training data and for the published baselines (Boltz-2, Protenix, Glide-SP, Gnina-Vina).

**Supplementary Figure 3:**
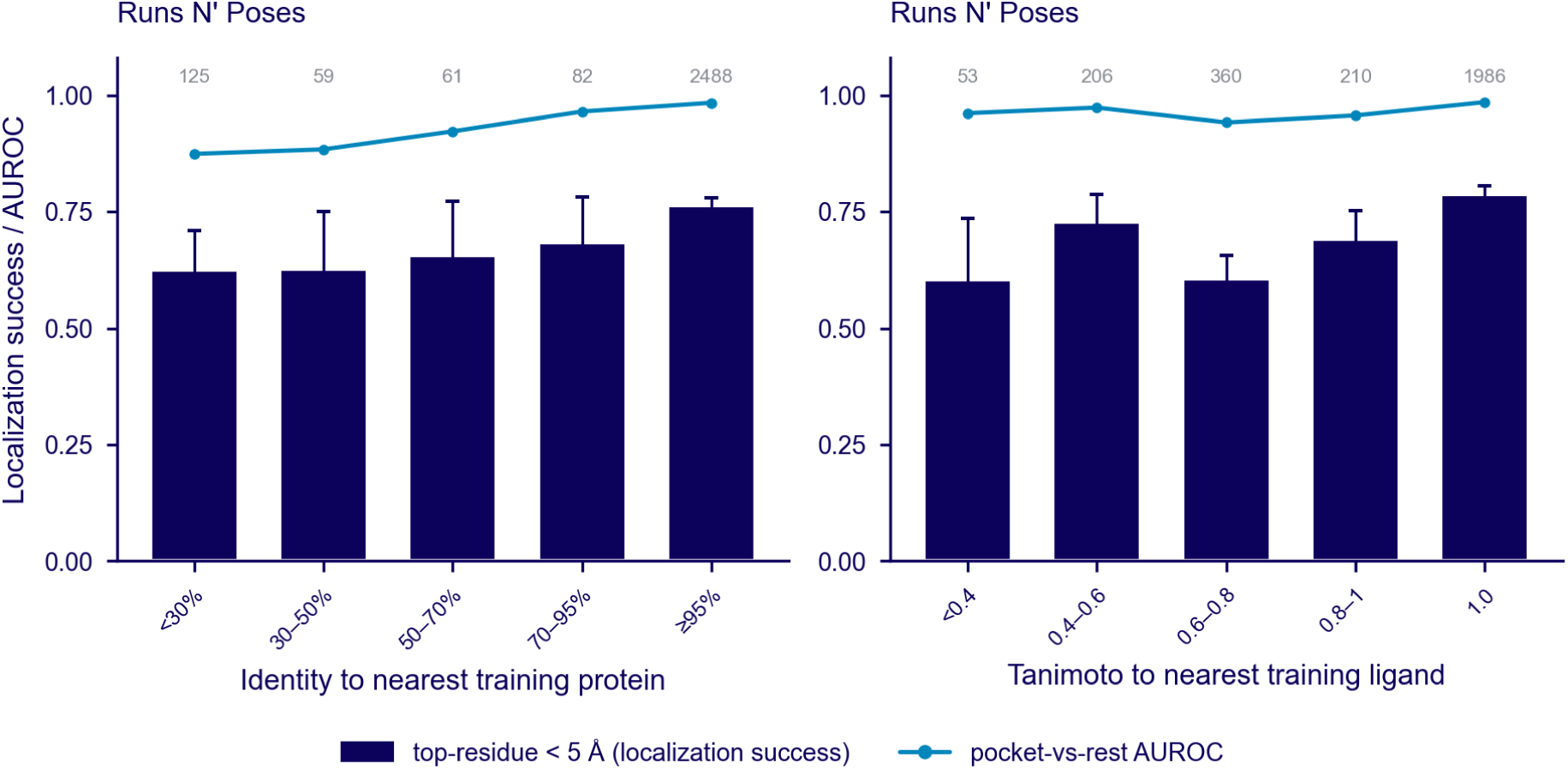
Localization on the Runs N’ Poses benchmark. [**23**] **as a function of similarity to Ptarmigan-1’s own training set, the companion to** Figure 2F. Bars, fraction of systems whose top-engagement residue lies within 5 Å of the ligand (exact-binomial 95% CI). Line, median pocket-vs-rest AUROC, per-bin counts above. (A) Binned by sequence identity to the nearest training protein. Lowest bin (identity < 30%, *n* = 125): top-residue recovery 62%, median pocket-vs-rest AUROC 0.87. (B) Binned by Morgan (ECFP4) Tanimoto to the nearest training ligand. Most-novel bin (Tanimoto < 0.4, *n* = 53): median pocket-vs-rest AUROC 0.96.

**Supplementary Figure 4:**
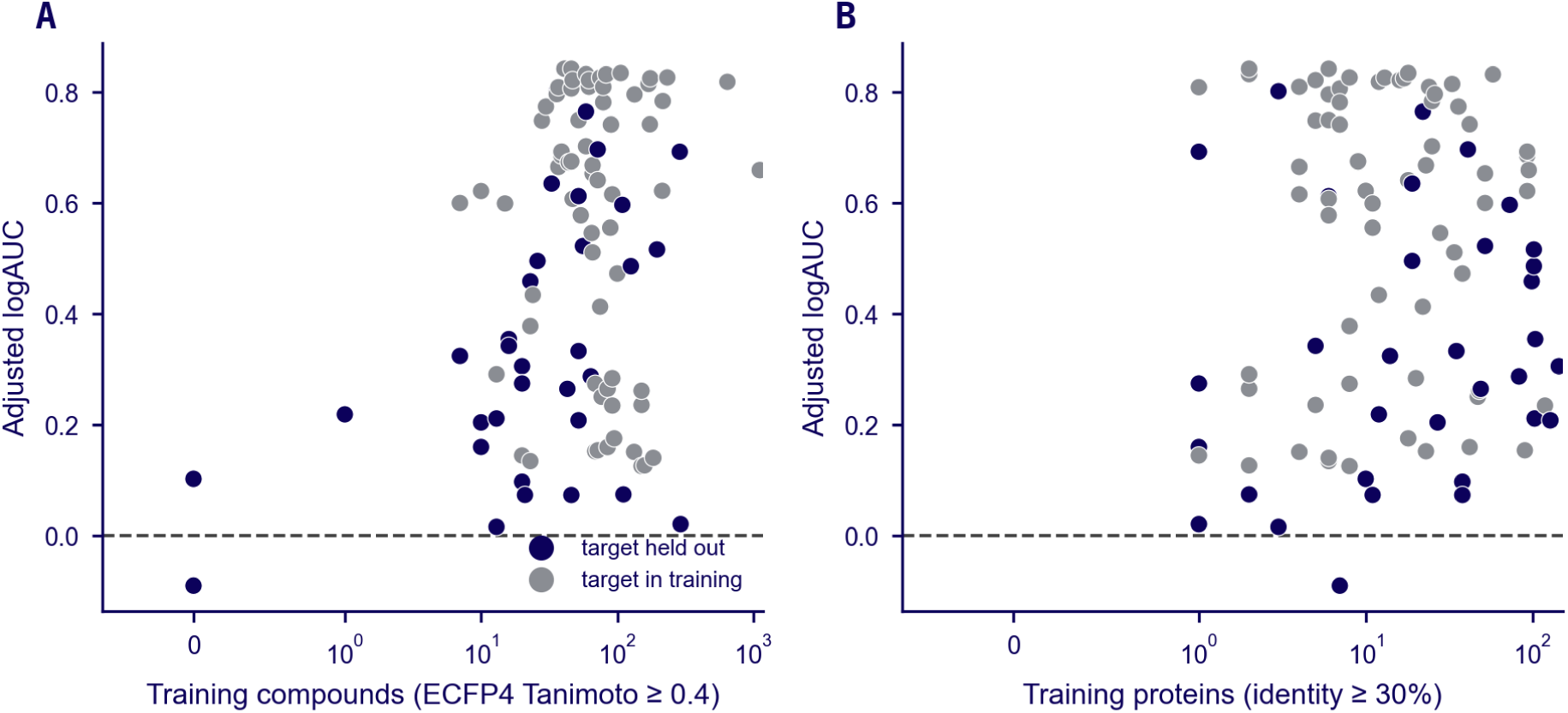
Per-target screening performance against chemical and protein coverage in training. For each of the 100 targets in our held-out screening benchmark we plot the per-target adjusted logAUC against a measure of how well the target is represented in training; points are colored by whether the target protein was withheld from training (dark; *n* = 33) or seen during training (grey; *n* = 67), the dashed line marks random performance (adjusted logAUC = 0), and both count axes are symmetric-logarithmic to accommodate targets with no training neighbor. (A) Chemical coverage, the number of distinct training compounds within a Morgan (ECFP4) Tanimoto of 0.4 of the target’s actives, taken as the median across actives and estimated from a uniform sample of the training library. (B) Protein coverage, the number of training proteins sharing at least 30% sequence identity with the target. Of the held-out targets, 32 of 33 score above random; the median adjusted logAUC is 0.31 for held-out targets and 0.65 for targets seen in training.

**Supplementary Figure 5:**
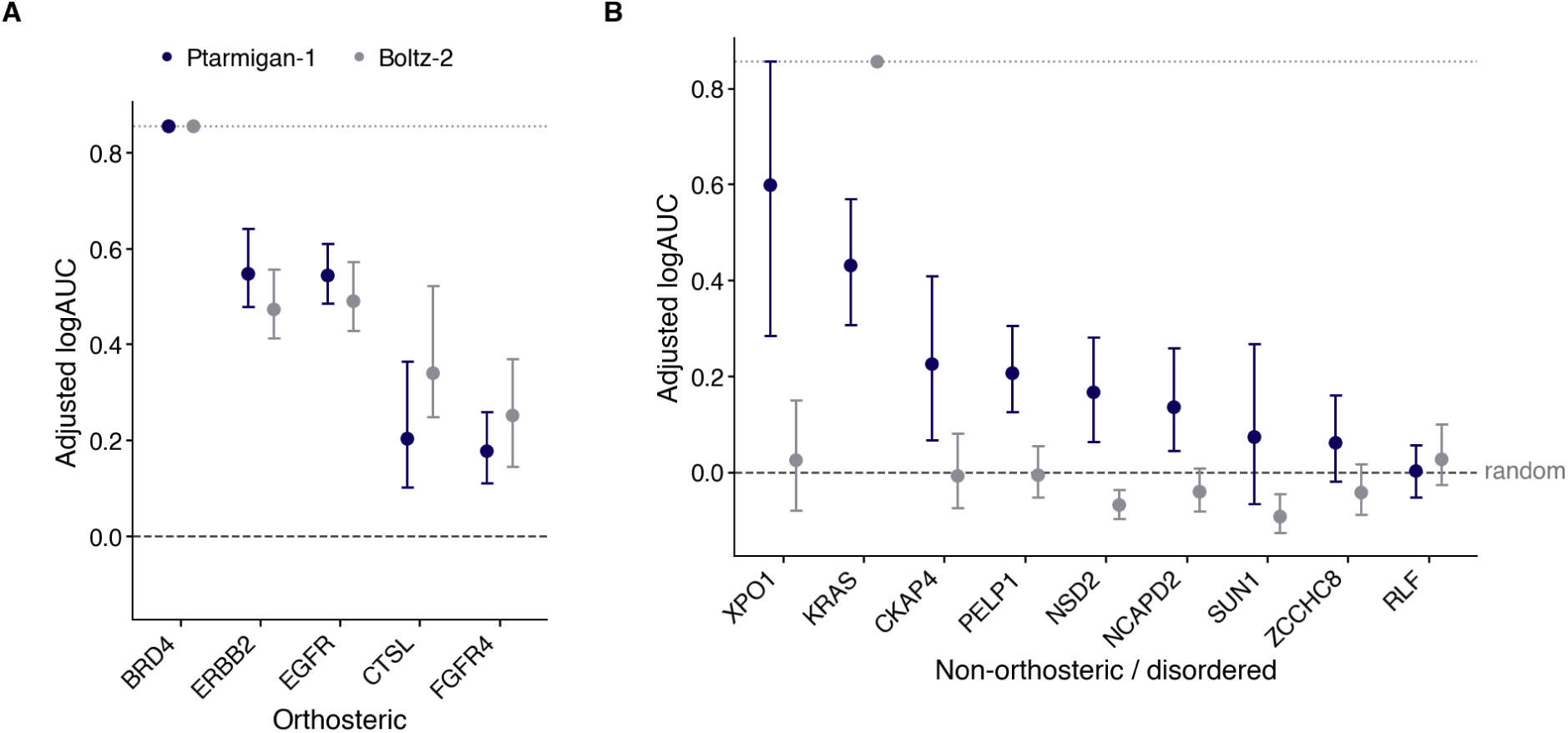
Per-target covalent virtual-screening performance, the detail behind the pocket-class summary in Table 2. Each point is a target’s adjusted logAUC (Mysinger–Shoichet, *λ* = 0.001) for Ptarmigan-1 (dark) and Boltz-2 (grey), with whiskers giving a bootstrap 95% confidence interval over the scored compounds. XPO1 here is the seven-target ChEMBL covalent benchmark (4 annotated actives), distinct from the larger, dedicated 37-SINE case-study screen in Figure 4E. Targets that reach the ceiling from perfect separation (BRD4, and Boltz-2 on KRAS) have a degenerate bootstrap interval and so sit on the dotted line without a whisker. Within each target both methods are scored on an identical set of compounds, retaining every active. (A) Covalent actives within the acetyl-lysine pocket of BRD4, the catalytic cysteine of cathepsin L (CTSL), and the ATP-site cysteines of the receptor tyrosine kinases EGFR, ERBB2, and FGFR4. (B) Covalent actives within non-orthosteric cryptic sites (KRAS, XPO1) and disordered protein regions.

**Supplementary Figure 6:**
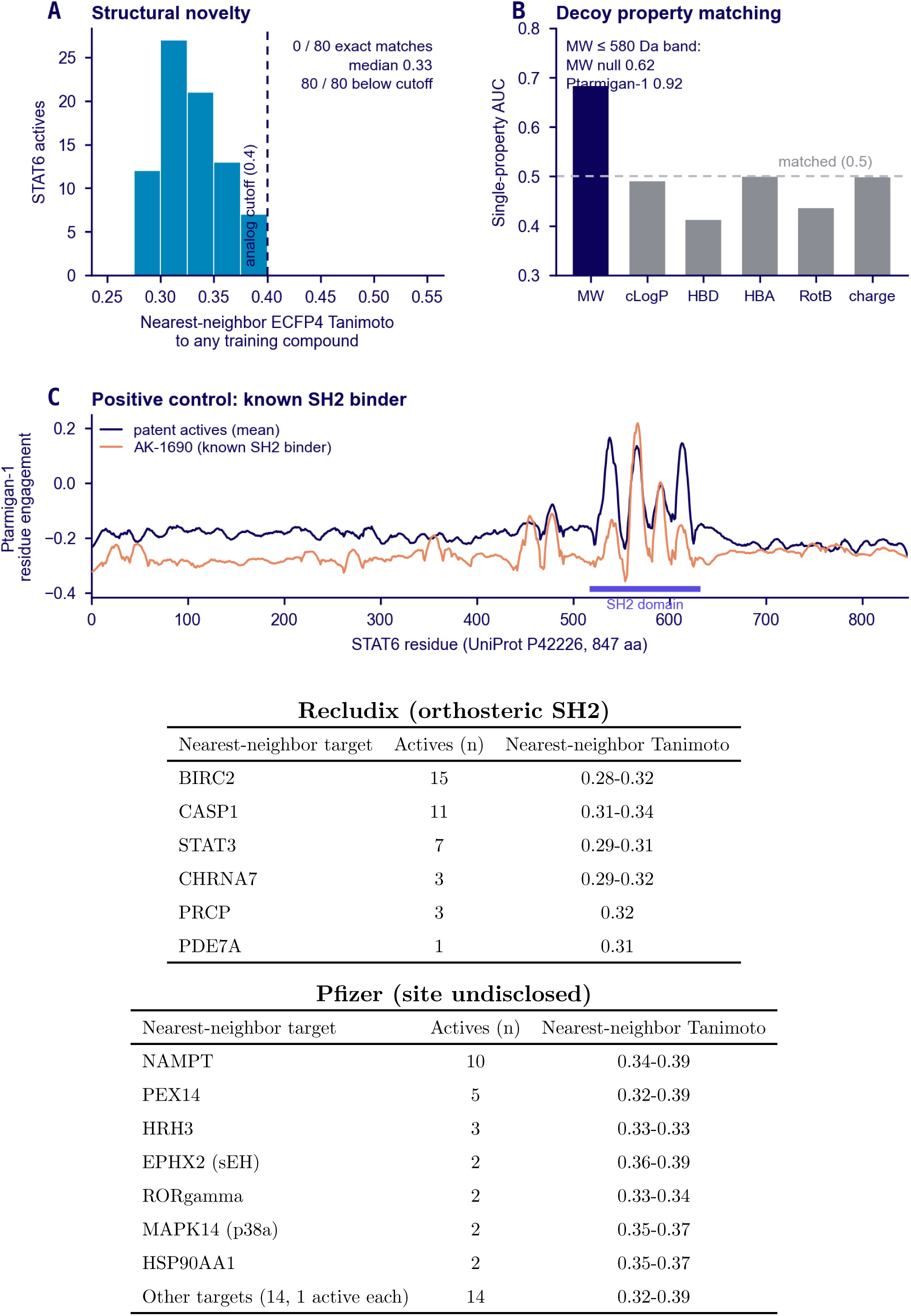
Structural-novelty, property-matching, and positive-control checks for the STAT6 case study. (A) The distribution of nearest-neighbor Morgan (ECFP4) Tanimoto to any compound in the full training corpus, over all 80 actives. There are zero exact structural matches, and the median nearest-neighbor is 0.33 (0.35 Pfizer, 0.31 Recludix). (B) The single-property ROC-AUC (active versus decoy) for six physicochemical properties over the matched library. (C) Ptarmigan-1’s smoothed per-residue engagement for AK-1690 (orange) overlaid on the patent-actives mean (dark). (D) What each series’ nearest training neighbors bind, grouped by target. The Recludix actives fall near a few chemotypes, including the STAT-family sibling STAT3 (7 of 40, Tanimoto ≤ 0.31); the Pfizer actives are spread across many unrelated targets. Neither series has STAT6 among its neighbors.

**Supplementary Table 1:** Ligand-conditioning controls for residue localization. Each readout of Figure 2 is recomputed on the same 337 drug-like PoseBusters complexes under three pairings. “Correct” scores each complex against its own crystallized ligand, reproducing the values in Figure 2D and Figure 2G. “Matched swap” scores it against a ligand belonging to a different protein, matched to the native ligand’s molecular weight (±50 Da), calculated logP (±1), and hydrogen-bond donor, acceptor, and rotatable-bond counts (±2 each), with no donor drawn from a complex sharing the query’s UniProt accession; five draws per complex are averaged before testing. “Agnostic panel” marginalizes per-residue engagement over a fixed 24-compound probe panel, excluding each complex’s own ligand. Δ is the paired per-complex difference from the correct-ligand arm, with a 95% percentile bootstrap interval over complexes; the Wilcoxon signed-rank p-value is below 10^−14^ for every readout (Methods). Binding-site recovery is the fraction of complexes whose single top-engagement residue lies within 5 Å of the ligand; precision at 5 and 10 is the fraction of the top-ranked 5 and 10 residues within that cutoff.

| Readout | Correct | Matched swap | $\Delta$ (95% CI) | Agnostic panel | $\Delta$ (95% CI) |
| --- | --- | --- | --- | --- | --- |
| Pocket-vs-rest AUROC | 0.968 | 0.916 | +0.053 (0.044–0.062) | 0.933 | +0.035 (0.028–0.043) |
| Binding-site recovery | 0.917 | 0.773 | +0.144 (0.107–0.179) | 0.807 | +0.110 (0.068–0.151) |
| Precision, top 5 | 0.922 | 0.797 | +0.126 (0.100–0.152) | 0.808 | +0.115 (0.085–0.144) |
| Precision, top 10 | 0.905 | 0.779 | +0.126 (0.104–0.148) | 0.797 | +0.108 (0.083–0.133) |

**Supplementary Table 2:**
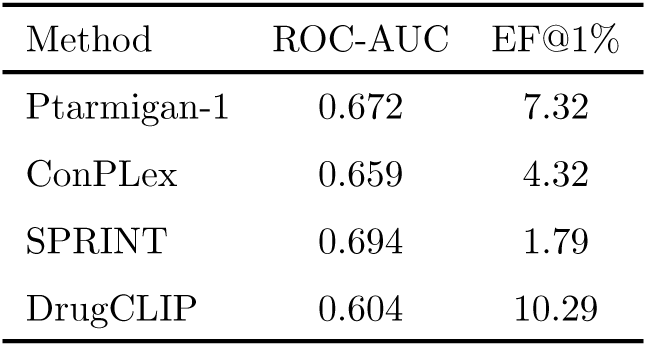
DrugCLIP [32] compared on LIT-PCBA, over the five well-folded target-sets that carry published structure-based baselines (estrogen-receptor-α agonist and antagonist, MAPK1, PPARG, and TP53). Entries are the mean ROC-AUC and enrichment at 1% (EF@1%). DrugCLIP published performance comes from its own evaluation pipeline and metric set (BEDROC at *α* = 80.5 rather than the *α* = 20 used throughout, and no adjusted logAUC). Ptarmigan-1, ConPLex, and SPRINT are scored on LIT-PCBA compounds, and DrugCLIP’s row is its own reported evaluation.

